# Conditions for optimal mechanical power generation by a bundle of growing actin filaments

**DOI:** 10.1101/344671

**Authors:** Jean-Louis Martiel, Alphée Michelot, Rajaa Boujema-Paterski, Laurent Blanchoin, Julien Berro

## Abstract

Bundles of actin filaments are central to a large variety of cellular structures, such as filopodia, stress fibers, cytokinetic rings or focal adhesions. The mechanical properties of these bundles are critical for proper force transmission and force bearing. Previous mathematical modeling efforts have focused on bundles’ rigidity and shape. However, it remains unknown how bundle length and thickness are controlled by external physical factors, and how the attachment of the bundle to a load affects its ability to transmit forces. In this paper, we present a biophysical model for dynamic bundles of actin filaments that takes into account individual filaments, their interaction with each other and with an external load. In combination with *in vitro* motility assays of beads coated with formins, our model allowed us to characterize conditions for bead movement and bundle buckling. From the deformation profiles, we determined key biophysical properties of tethered actin bundles, such as their rigidity and filament density. Our model also demonstrated that filaments undulate under lateral constraints applied by external forces or by neighboring filaments of the bundle. Last, our model allowed us to identify optimum conditions in filament density and barbed end tethering to the load for a maximal yield of mechanical power by a dynamic actin bundle.

## Introduction

Bundles of actin filaments are involved in a large variety of cellular structures, such as filopodia (1), the cytokinetic ring (2, 3), auditory hair cells (4), actin cables (5, 6), stress fibers (7, 8), focal adhesions (9), adherens junction (10), and are hijacked by some pathogens for their propulsion within and between host cells (11). Bundles also determine the shape of macroscopic structures like Drosophila bristles (12) and are the core of microvilli (13, 14).

In motile cells, bundles of actin filaments can develop large amounts of force that deform the cell membrane at the leading edge (13, 15, 16), and are used to generate tension in stress fibers (17). Filopodial bundles are created by formin- and Ena/Vasp- mediated assembly of parallel actin filaments. In the lamella, actin filaments from the lamellipodium are condensed into bundles under the action of the retrograde flow and motor proteins. Through these bundles, cells optimize and adapt their response to mechanical stress and disassemble the actin skeleton at their trailing edge (18–20).

Experimental evidence has shown that the force exerted by an untethered bundle of actin filaments against a wall is virtually equal to the force exerted by a single filament (16). This surprising result was attributed to a rapid switch between the leading filaments at the tip of the bundle, such that only one filament at a time contacts the load. We speculate that this result would have been different if all filaments were tethered to the wall. In addition to filament-load interactions, another limiting factor for force production is filament buckling (20–22). Buckling occurs when the force exerted between the ends of the filament reaches a critical value, which depends on the mechanical properties of the filament and the attachment conditions of its ends (23, 24). When buckling occurs the force produced by a filament (or a bundle) vanishes.

Previous theoretical and *in vitro* biophysical studies have determined the mechanical properties of bundles of actin filaments (15, 25, 26), and determined the effects of crosslinkers (27), motor proteins (21, 22, 28) or active transport (29). These studies essentially focused on the effect of bundle rigidity on its stability and shape, and how filament dynamics and membrane tension control the bundle length. However, it remains unknown how bundle length and thickness are controlled by external physical factors, such as bulk viscosity or the spatial constraints applied on the bundle by its attachment to a load and to other filaments, or by other cytoplasmic structures.

In this paper, we present a biophysical model for bundles of dynamic actin filaments taking into account individual filaments and their interactions with a load and with other filaments of the bundle. The model is supported by experimental *in vitro* reconstitution of bead motility powered by bundles of actin filaments, which are nucleated on the bead surface by formins. Here, formins also play the role of tethers that maintain filament barbed ends on the bead. Using our experimental data we quantified the bead movement and correlated it with the elongation and deformation of filament bundles. This allowed us to determine bundle’s rigidity and average number of filaments. Thus, using these two experimentally-measured constraints, we propose an original, simple and robust model for the movement and deformation of actin filaments that predicts how filament density and tethering of to the load tune the mechanical power produced by dynamic actin filament bundles.

## Materials and Methods

### Experimental data

#### Motility assay

Experiments performed in this study were carried out according to the procedure described in (30). In brief, beads grafted with the FH1-FH2 domains of mDia1 were mixed with a motility medium containing 8 µM F-actin, 4 µM profilin, and 10 µM human cofilin in X buffer (10 mM HEPES pH 7.8, 0.1 M KCl, 1 mM MgCl2, 1 mM ATP, and 0.1 mM CaCl2), supplemented with 1% BSA, 0.2% methylcellulose, 3 mM DTT, 1.8 mM ATP, and 0.1 mM DABCO.

#### Microscopy

Motility assays were acquired with a Zeiss Axioplan microscope (Jena, Germany) equipped with a 63x/1.5NA Plan-APOCHROMAT objective lens, a Hamamatsu ORCA CCD camera (Hamamatsu Photonics Deutschland GmbH) and Metavue version 6.2r6 (Universal Imaging, Media, PA).

Buckles of filaments were imaged using Total Internal Reflection Microscopy (TIRFM). Glass flow cells were cleaned and prepared according to (31). Rhodamine-actin and Alexa-532-labeled actin-filament polymerization was observed and acquired as specified in (32).

### Theory and simulations

We present below the 3D model for elastic filaments (Eqs. 1-2). In the supplementary material, we simplify this 3D model to develop a general method for the analysis of filaments or bundles of filaments considered as elastic structures in 2D (Eqs. S1-S12). We estimate the relative importance of the polymerization and drag force (Eqs. S13-S15) involved in the bead movement (Eqs. S16-S20). Finally, we use the 3D model (Eqs. 1-2) to analyze the role of attachment and compressive forces due to filament packing to determine the mechanical power produced by individual filaments (Eqs. S21-S36).

#### Mechanical equilibrium equations for an actin filament

We developed a model for the mechanical equilibrium of elastic filaments subjected to external forces and constraints based on a model we previously developed (33). In this section, we present the definitions of variables and the equations for force and moment balance. The orientation of the filament cross-section at any point along the filament is given by a set of three unit vectors (**d**_1_, **d**_2_, **d**_3_) which define the material frame associated with position **r**(s) (Fig. S1A). The filament bending or twisting strain **κ**, the density of force **f,** and moment **m** vectors are defined in the material frame as **κ** = *κ*_1_**d**_1_ + *κ*_2_**d**_2_ + *κ*_3_**d**_3_, **f** = *f*_1_**d** + *f*_2_**d**_2_ + *f*_3_**d**_3_, **m** = *m*_1_**d**_1_ + *m*_2_**d**_2_ + *m***d**_3_. The moment is proportional to the filament bending strains **m** = *B***κ** where *B* is the bending rigidities diagonal matrix diag(*C_B_*, *C_B_*, *C_T_*). The balance of force and moment reads

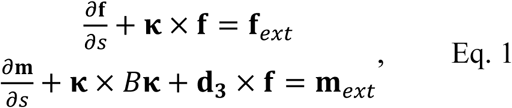

where **f**_ext_ and **m**_ext_ are the external force and moment densities applied to the filament at position *s*. All filament-filament or filament-medium interactions are modeled by adapting the expression for **f**_ext_ in Eq. 1. The unit tangent vector, denoted **d**_3_, is given by

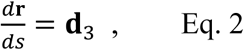

where **r** is a point along the filament (Fig. S1A). Since filaments move at low Reynolds number (∼ 10^−5^), all force and moment terms proportional to linear or angular acceleration were eliminated from Eq. 1 (33). The components of the material frame (**d**_1_, **d**_2_, **d**_3_) vectors are parametrized by Euler parameters (unit quaternions), which couple the change of the cross section orientation to the force-moment applied at *s*.

**Figure 1.**
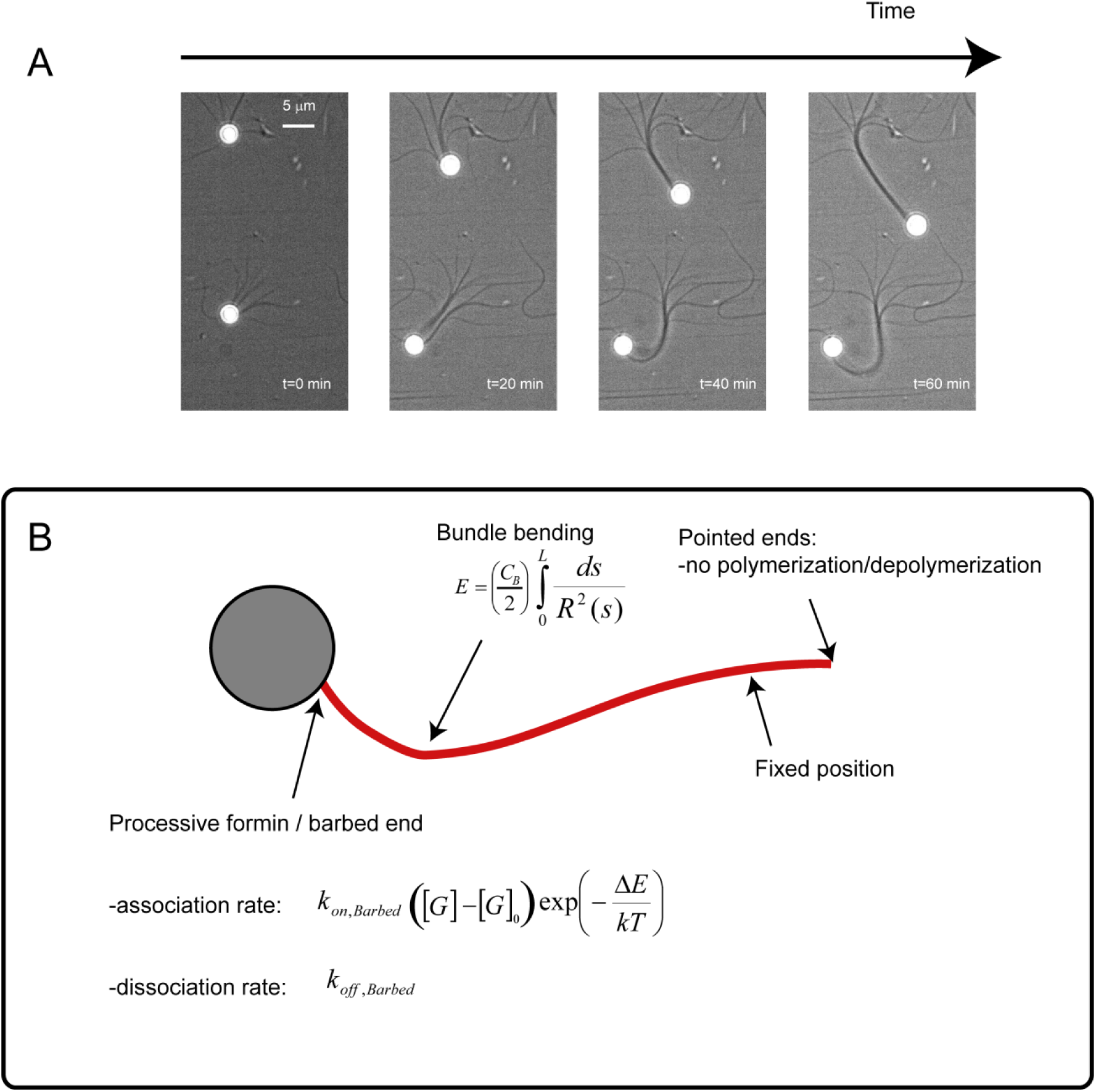
Bead motility mediated by bundles of actin filaments nucleated by processive formins (mDia1). <u>Panel A:</u> Beads coated with formins mDia1 (20 to 160 formins/bead) are added to a F-actin motility medium (8 µM F-actin, 4 µM profilin and 10 µM ADF/cofilin.) Actin filaments are arrayed in parallel bundles with their barbed at the bead surface. In some cases, the force developed by actin polymerization propels the bead at constant velocity (∼ 0.25-0.27 µm.min^−1^) for up to one hour (bundle and bead at the top). In other cases, the driving force fails to move the bead after a certain time of steady displacement (e.g. 30 min for the bead at the bottom) while polymerization continues, as demonstrated by bundle deformation. <u>Panel B:</u> Schematic of the mechanical model for bead motility and bundle deformations. We combine the mechanics of bundle with elongation at the barbed end. The polymerization rate is corrected by a term depending on AE, the mechanical work necessary to insert a monomer.

#### Microscopic model for buckling

We used the 3D model for actin filaments (Eqs. 1-2) to analyze the role of 1) attachment conditions (tethered vs. non-tethered filaments) and 2) filament density in the emergence of an optimum for mechanical power production. For both attachment conditions, the actual filament elongation rate depends on the force component *FN* normal to the bead surface through the empirical relation

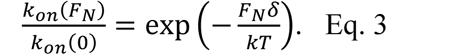

In the simulations, we neglected the depolymerization at the barbed end.

##### Boundary conditions for tethered filaments

The boundary condition at the pointed end specifies the position **r** of the filament and its orientation **q** (a total of 3+4 conditions)

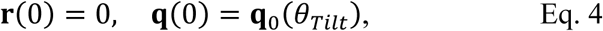

where **q**_0_ (*θ_Tilt_*) gives the direction of the filament at the pointed end. At the barbed end, we impose zero bending-twisting strains (a total of 3 conditions)

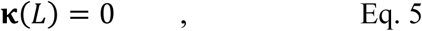

and the position of the barbed end is fixed in space

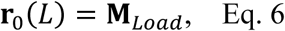

where **M***_Load_* is a point at the load surface.

##### Boundary conditions for non-tethered filaments

The boundary conditions at the pointed end are similar to the boundary conditions for tethered filaments (Eqs. 4 and 5). At the barbed end, only the horizontal position of the barbed end is constrained (one condition)

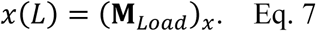

We complete the set of boundary conditions for non-tethered filament by imposing zero tangential force at the barbed end to model the absence filament slippage on the bead (2 conditions)

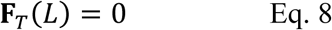

Finally, for both types of attachment, we end up with 13 boundary relations which are used to simulate the equilibrium of filaments with Eqs. 1-2. The constraints exerted on a filament due to the environment are imposed via the right hand side of Eq. 1a, which reads

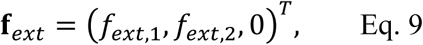

where *f*_ext,12_ are, respectively, the components of the force density along the first and second director vectors. The expression of **f_ext_** on the surrounding filament density and other geometric factors is given in the **Result** section and by Fig. S10.

## Results

### Buckling of actin filament bundles limits the motility of formin-grafted beads

When placed in a motility medium containing actin monomers, profilin and cofilin (30, 34), beads coated with formin mDia1 nucleate actin filaments at their surface (Fig. 1A). These elongating filaments rapidly form long bundles of actin filaments, which propel the bead at a constant velocity about 0.25 µm.min^−1^ for a duration that varied from a few minutes to over one hour, when the experiment was stopped (Fig. S5). In most cases, beads eventually stalled and remained permanently stuck to the coverslip (Figs. 1A and S2). During the motility phase, bead trajectories were rectilinear, their bundle remained straight (Fig.1A), and the bead velocity and the bundle elongation rate were identical (Fig. S2). After beads stalled, bundles continued growing at the same elongation rate while deforming to eventually form a large buckle (Fig. 2).

**Figure 2.**
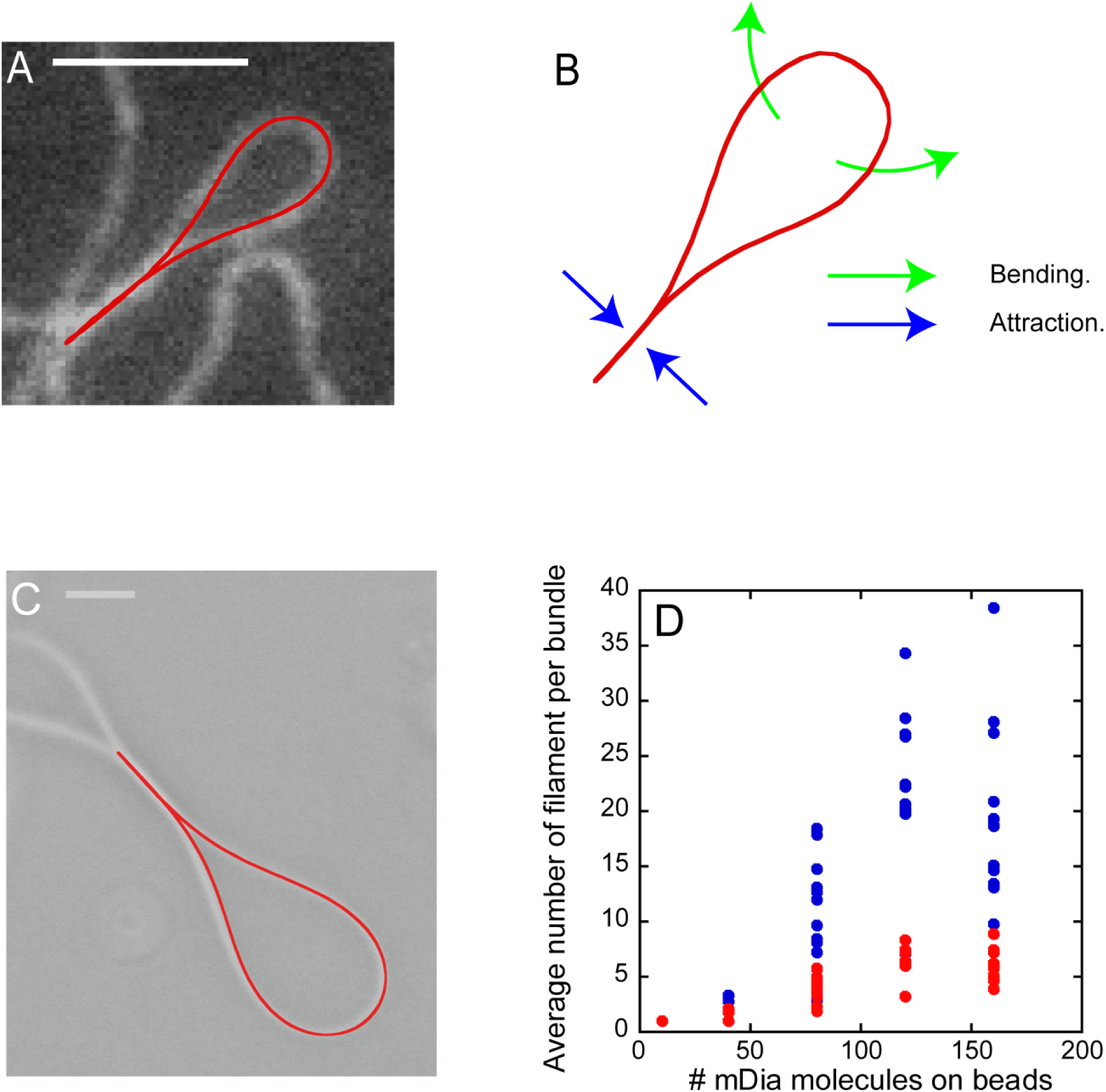
Formation of loops by individual filaments and bundles of filaments. <u>Panel A:</u> Single filament can form a loop, which can be fitted with our 2D model for elastic filaments. Bar: 5 µm. <u>Panel B:</u> Cartoon showing how attractive forces along the loop stem (blue arrows) balance the elastic forces caused by filament bending (green arrows). <u>Panel C:</u> Superposition of a loop formed by a bundle of filaments and the associated solution from our 2D model. Bar: 5 µm. <u>Panel D:</u> Number of actin filaments in bundles estimated from rigidity measurement assuming sliding conditions (β=1, blue dots) or no sliding conditions (β=2, red dots).

What is the cause of bead movement and why do beads stall? Since filaments in bundles grow at the same rate when beads are moving and when beads are stalled, we deduced that actin monomer concentration and viscosity of the motility medium remained constant (Fig. S5). In consequence, these two factors cannot account for the changes of the bead velocity. We hypothesized that the variety of motility behavior observed experimentally is due to the variations in the mechanical properties of the bundle of actin filaments over time.

### Determination of bundle rigidity from loop geometries

To test our hypothesis, we took advantage of loops formed by long filaments (Fig. 2A) and bundles of filaments (Fig. 2C) at the end of our experiments. We assumed that these loops are elastic structures at equilibrium, where the elastic bending force of the filament (or bundle) which tends to open the loop, and the attractive depletion forces that keep the stem of the loop closed, balanced each other (Fig. 2B, Eqs S1-S2, and *Supplemental text, section A*).

The persistence length of bundles can be directly deduced by fitting the shape of the loops to our model (Eqs. S5-S6 and Figs. S2D, and S6). As expected, the apparent persistence length of bundles increased with the number of formin molecules bound to the bead and the nature of filament-filament interactions within the bundle (Figs. 2D and S7). However, our results show that the number of filaments is significantly smaller than the number of formins on the bead estimated experimentally, and plateaus for beads with more than ∼100 formins. This discrepancy can be partly explained by the fact that not all the adsorbed formins are functional. This argument can also explain the large variability in the number of filaments nucleated by a bead (Figs. 2D and S7). In addition, as we show below, the number of filaments attached to the bead is limited by mechanical stress applied by elongating filaments on the formins. Overall, the variability in the number of filaments in bundles likely explains the diversity of bead trajectories or loops observed in the same experimental conditions (Figs. 1A and S2).

### Bead stalling as a transition between polymerization-dominated to elasticity-dominated regimes

To test whether the difference in the mechanical properties of the bundle can explain our experimental results, we developed a model for bundle mechanics and bead movement (Fig. 1B), where the bundle is modeled as a single elastic rod with bending rigidity *L_P_* and contour length *L (Supplemental data, section B)*. The bundle elongates from its barbed end which is attached to the bead via formins. The other end of the bundle is assumed to have a fixed position and orientation on the coverslip, as experimentally observed. The transition between bead movement and bundle deformation depends on the balance between two antagonistic forces, namely 1) the force produced by actin polymerization and 2) the viscous drag due to the viscosity of the medium on the bead (Fig. 3A). The physical origin of these forces and their order of magnitude are given in Table 2. If the force required to deform a bundle is higher than the drag force, the bead is steadily pushed forward while the bundle remains straight (Fig. 3A, left). Conversely, when the viscous drag is larger than the critical buckling force of the bundle, the bead stalls, and the force generated by the elongation deforms the bundle (Fig. 3A, right). This critical buckling force scales as the reciprocal of the square bundle length (Eqs. S13). Therefore, in conditions of constant elongation, the bundle length will eventually meet the condition for buckling, and the exact time for this transition to occur depends on the bundle rigidity or, equivalently, the number of filaments in the bundle (Eq. S11).

**Figure 3.**
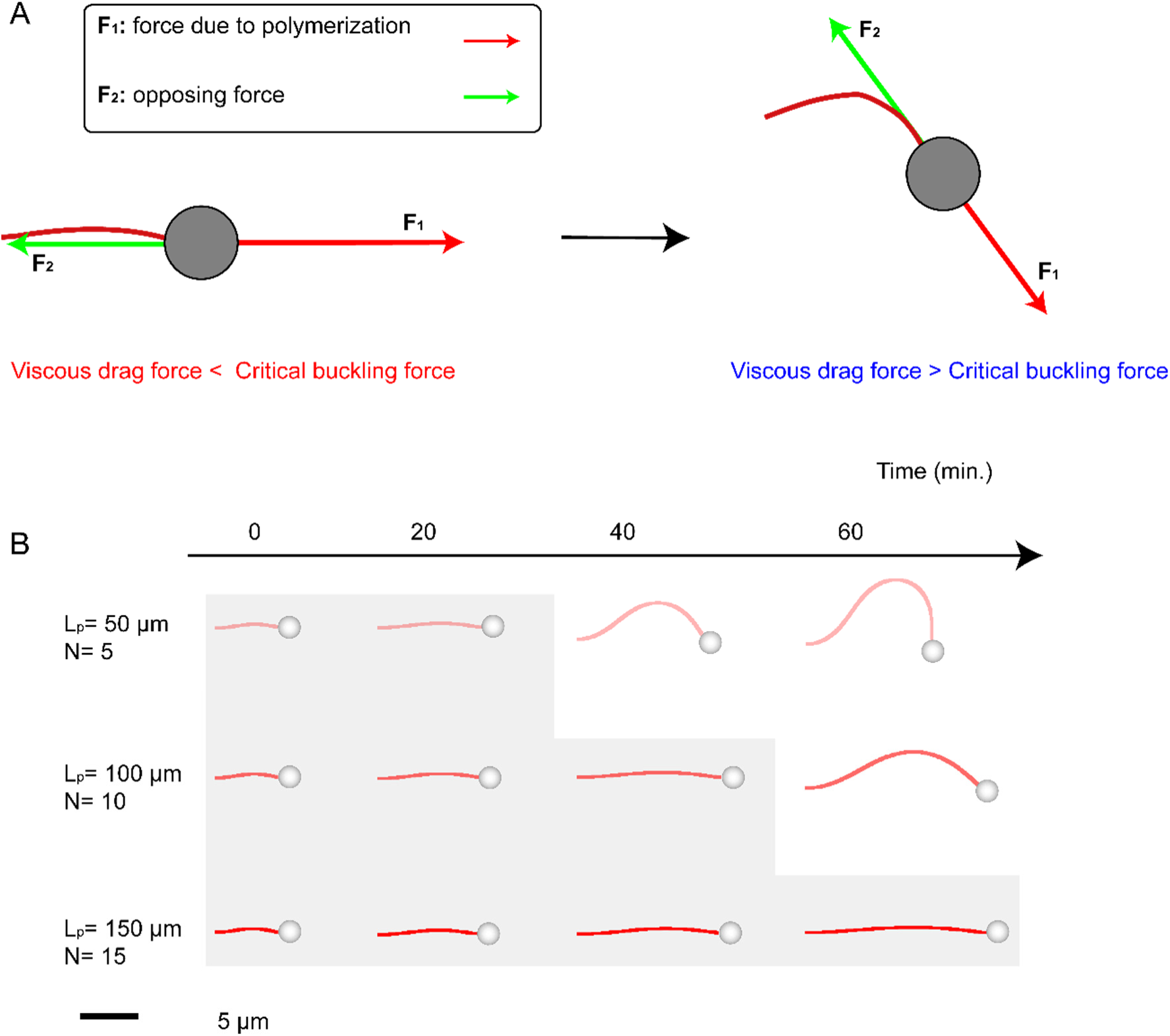
Motility and bundle deformation. <u>Panel A:</u> The balance between polymerization and drag forces on the bead controls the transition from bead movement to bundle deformation. <u>Panel</u> B: The bundle rigidity controls the transition between bead movement and bundle deformation. Shaded areas give conditions for which bundles generate enough force to propel the bead.

Our mathematical model for bead movement (Eqs. S16-20) reproduces the transitions between both kinds of movement (Figs. 3B and S5, left column) and shows the dependence of this transition on bundle rigidity *L*_P_. Bundles with relatively low persistence lengths (e.g. *L*_p_=50µm) cannot propel the bead throughout the medium over a long period of time (Fig. 3B). Once the critical buckling condition is reached, i.e. when the bundle reaches the critical length (Eq. S15), the bead stops while the bundle continues its growth. This situation is persistent over time, since the elastic force produced by the bent bundle diminishes with the bundle length as

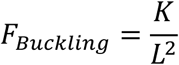

where *K* is a constant (see Eq. S13). In the case of high bundle rigidity (e.g *L*_p_=150µm), the bundle stays straight for about one hour during which the bead moves at a constant velocity without bending the bundle (Figs. 3B and S5, bottom right). The shape of the bundle when the bead is stalled (Fig. 3B) is in very good agreement with experimental data (Fig 1A).

Our model also allowed us to compute the critical length (*LCrit*, Eq. S15) at which a bundle undergoes the transition from force transmission to the bead to bundle deformation. This critical length scales as the square root of the persistence length (Fig. 4A, red curve), since the ratio 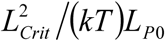 is constant (Eq. S15). This simple equation allows us to draw a boundary between bead motility and bundle deformation (Fig. 4A). During a typical experiment, the persistence length of the bundle for a given bead remains constant while its length increases. Therefore, it is represented by a trajectory along a vertical arrow in the phase diagram (Fig. 4A). The bead starts with a short bundle (under the phase transition boundary), then elongates while pushing the bead forward, until the bundle eventually reaches the critical length. Above this critical value, the bundle bends and the bead stalls (Figs. 1A and 4A, dashed arrow). Increasing viscosity of the medium increases the resistance to movement and shifts the phase transition boundary downwards in the phase space without altering its shape (Fig. 4B, Eq. S15).

**Figure 4.**
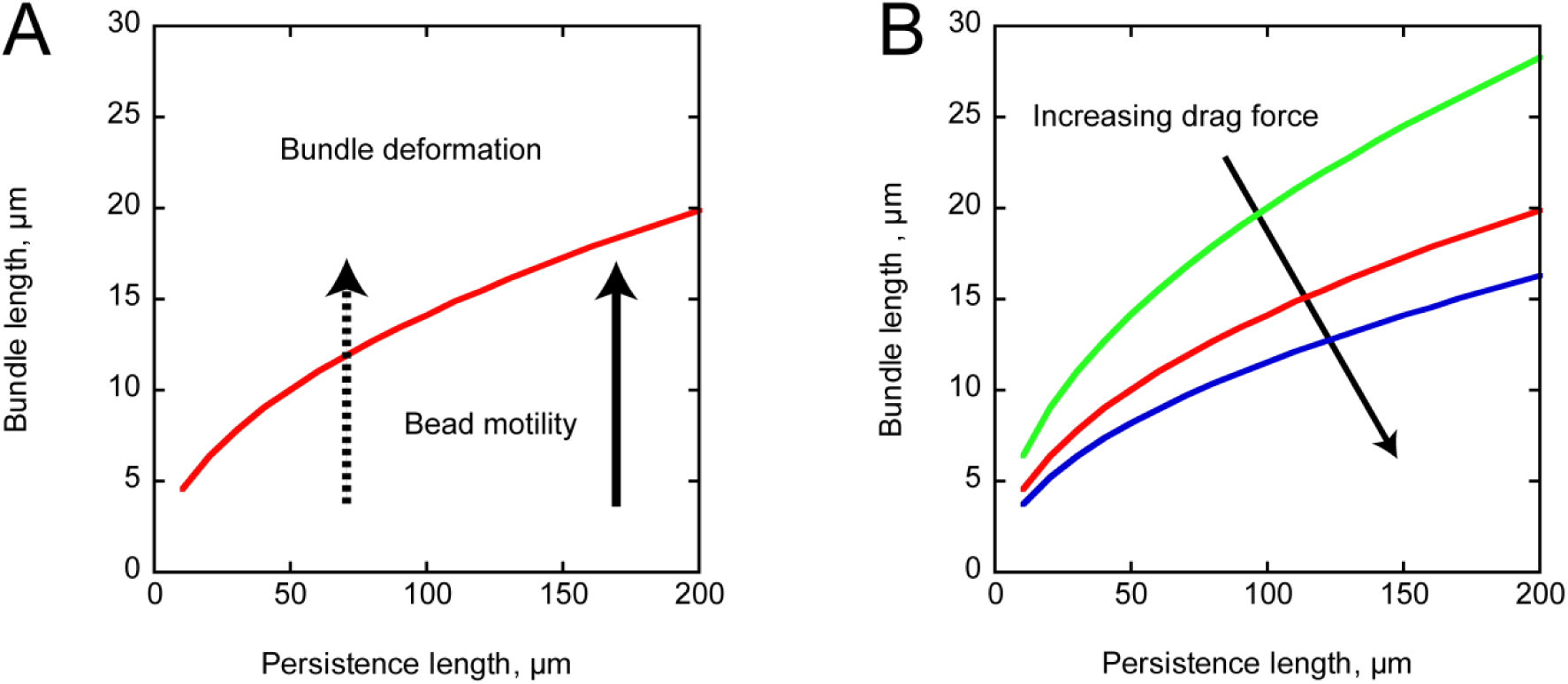
Phase diagram for bead movement and bundle deformation. <u>Panel A:</u> Transition between bead motility and bundle deformation in the parameter space (bundle length vs bundle rigidity) corresponding to simulations in Fig. 3. The dashed and solid arrows illustrates the typical trajectory in the phase diagram space associated with, respectively, a stalling bead (Fig. 1A, bottom, Figs. S5A and S5C) and a moving bead (Fig 1A, top, Figs. S5B and S5D). <u>Panel B:</u> Increasing the viscous drag force controls the position of the transition curve separating motility and bundle deformation.

### Emergence of optimal conditions for mechanical power yield

*In vitro*, the force production is limited by the elasticity of the bundle (Figs. 1 and 4), which is under the control of the filament interactions in the bundle. To better understand the relationship between bundle organization and the resulting mechanical work, we used the 3D model (Eqs. 1-9) to simulate a population of short actin filaments whose pointed ends are immobile and oriented with *θ_Tilt_*, the angle between the filament and the horizontal axis (Fig. 5).

**Figure 5.**
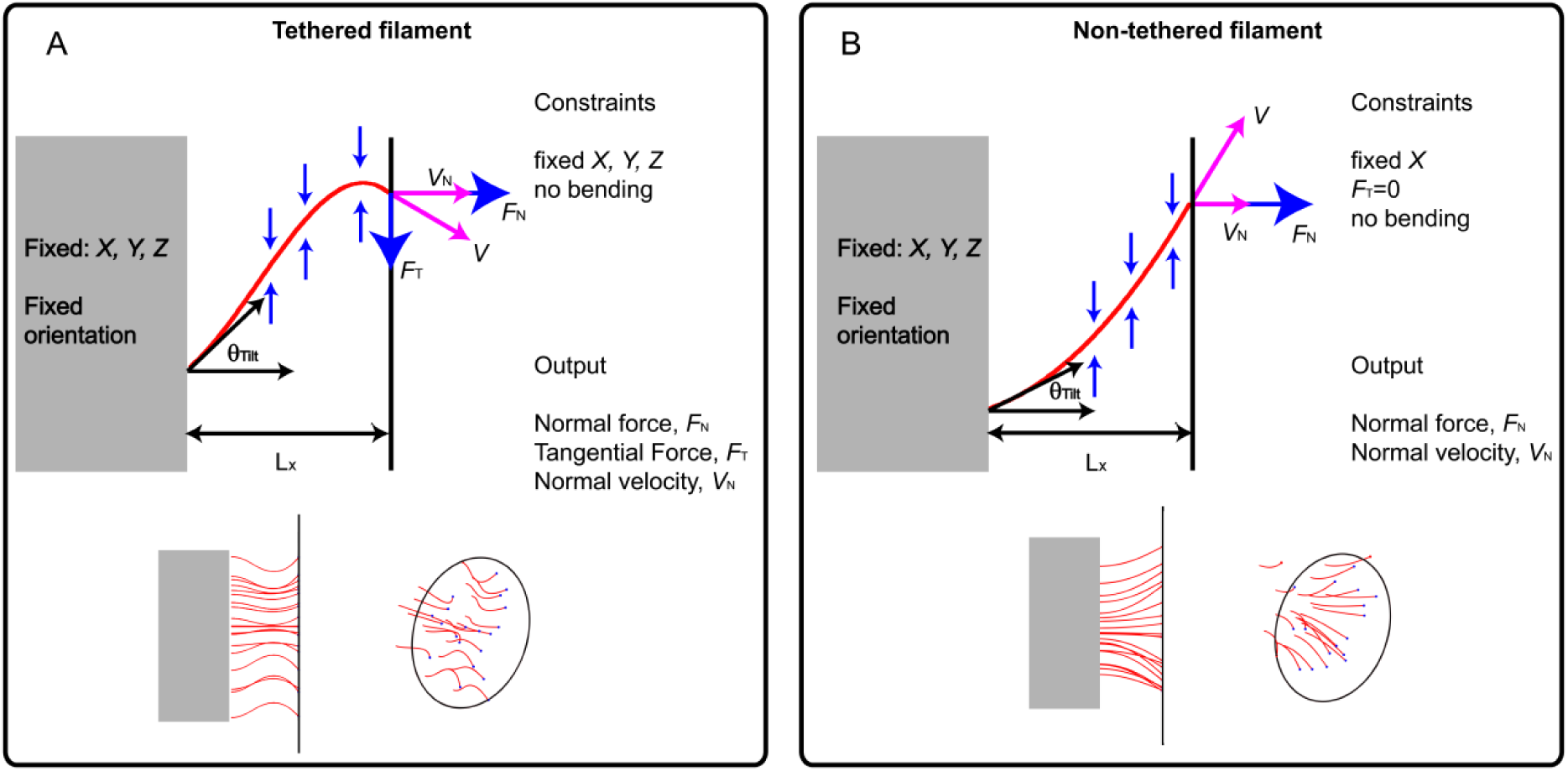
Microscopic 3D model for buckling of filaments in a bundle. The barbed ends of a short actin filament within a bundle (2 µm) pushes against a load (vertical line on the right) while its pointed ends is maintained at a fixed position and orientation (grey box on the left). Lateral interactions between filaments result into elastic compressive forces on individual filaments (vertical blue arrows) preventing the formation of a large buckle. <u>Panel A:</u> Tethered barbed ends exert normal and tangential forces FN and FT pushing the load at velocity V (magenta arrow). Bottom: side and perspective drawing of buckled filaments (red) tethered (blue points) to a load. <u>Panel B:</u> Non-tethered barbed ends exert a normal force FN only, with zero tangential component. The bottom cartoon shows the non-tethered filaments pushing against a load.

The barbed ends of all filaments are either bound to the bead by a link (e.g. a formin) (Fig. 5A) or freely pushing against the load (Fig. 5B), which, for simplicity, is modeled as a flat surface. Note that tethered filaments exert both a normal and a tangential force (Fig. 5A), whereas non-tethered filaments exert a normal force only (Fig. 5B). In addition, filaments are subjected to lateral constraints resulting from other surrounding filaments in the bundle (Fig. 5, small blue arrows). The lateral constraints prevent the buckling of individual filaments in the bundle and coerce the filament to stay in the bundle.

### Filament confinement changes the waviness of filaments and the force they exert along the end-to-end axis

We assumed that the lateral constraints in dense networks coerce the filament to remain in a cylinder whose radius decreases with the filament density (Fig. 6, top panel). To get a semi-analytical expression of the filament shape under compressive lateral load mimicking the presence of surrounding filaments, we simplified Eqs. 1-9 into a 2D model (Supplemental text, section C).

**Figure 6.**
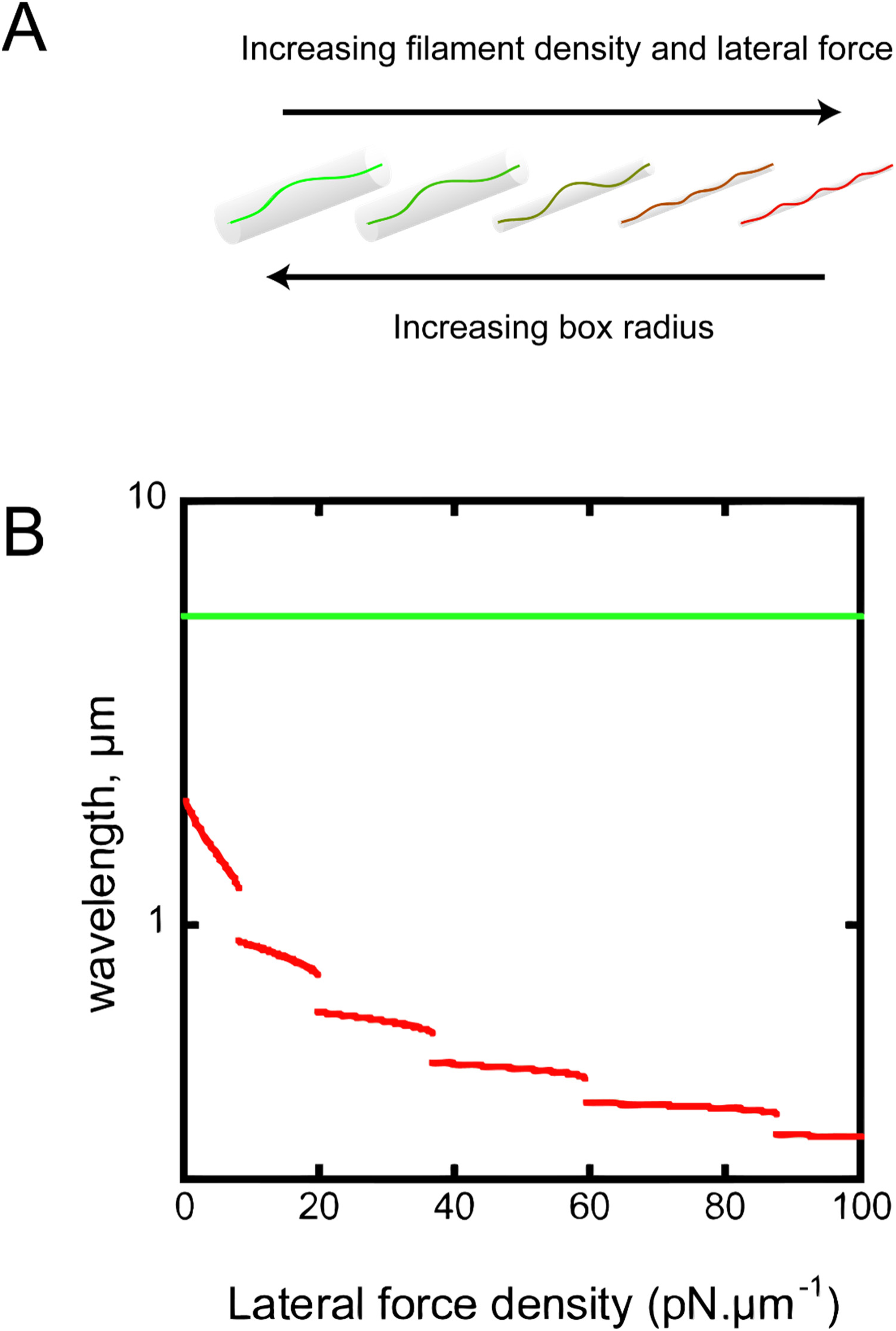
Filament shape is controlled by lateral constraints. *<u>Panel A</u>*: Shapes of a typical tethered filaments for increasing lateral force density (0, 20, 40, 60 and 80 pNµm^−1^ from left to right) which coerces filaments into cylindrical boxes with diminishing radius. *<u>Panel B</u>*: The wavelength λ, associated with ground state solutions (solutions of minimal elastic energy) are extracted from Fig. S9. Red (resp. green) curve corresponds to tethered (resp. non-tethered) conditions. All curves are obtained for a 2-µm filament that is moderately compressed along its end-to-end axis (end-to-end distance: 1.6 µm).

When filaments are isolated, they form a single buckle. However, when they are constrained to grow in a cylinder (which mimics the presence of surrounding filaments), they adopt a wavy shape (Figs. 6A and S11). These filament undulations are described by a single parameter, *λ*, the *wavelength* of the filament. Here, the *wavelength* is defined as the distance between two successive bumps (Figs. 6A, S8A and S11). The implicit equations for the filament wavelength (Eqs. S28 and S34) admit an infinity of solutions (Fig. S9), but only solutions of minimal energy (the so-called ground state) associated with the maximal wavelength can be observed. The equation also shows that high lateral force density correlates with highly wavy filaments, i.e. with a short *wavelength* (see Eq. S24).

The wavelength of the ground state solution for tethered filaments is large (∼5µm) and independent from the lateral compression exerted on the filament (Fig. 6, bottom panel, green curves). Therefore, the force *f*_x_ exerted by non-tethered filaments (Eq. S24) is always low and does not depend on the presence of surrounding filaments. In marked contrast, the wavelength of ground state solutions corresponding to tethered filaments decreases dramatically with increasing lateral constraints (Fig. 6, bottom panel, red curve). In consequence, tethered conditions favor highly tortuous filament configuration with high magnitude longitudinal force *f*_x_ (Eq. S3). Note also that the filament wavelength has jump transitions (Fig. 6 bottom panel) as the lateral force is increased continuously. Because of the boundary conditions (Eqs. S26, S28, S30 and S34), the number of waves along a filament is always an integer (resonance condition, see Fig. S8A). Therefore, by increasing the lateral force *f_x_*, one diminishes the wavelength λ (see Eq. S24). In consequence, the resonance condition imposes that the change in λ should be at least half an undulation (see Fig. S8B); hence the discontinuities observed in Fig. 6.

Using the full 3D model (Eqs. 1-2), we show that tethered filaments generate high magnitude force at the filament end whereas non-tethered filaments exert very little force (Figs. 7A and B). In addition to filament density, the orientation of the filament relatively to their attachment sites towards their pointed end, *θ*_Tilt_, plays also a significant role in the level of the force reached at large density (Figs. 7A and B).

**Figure 7.**
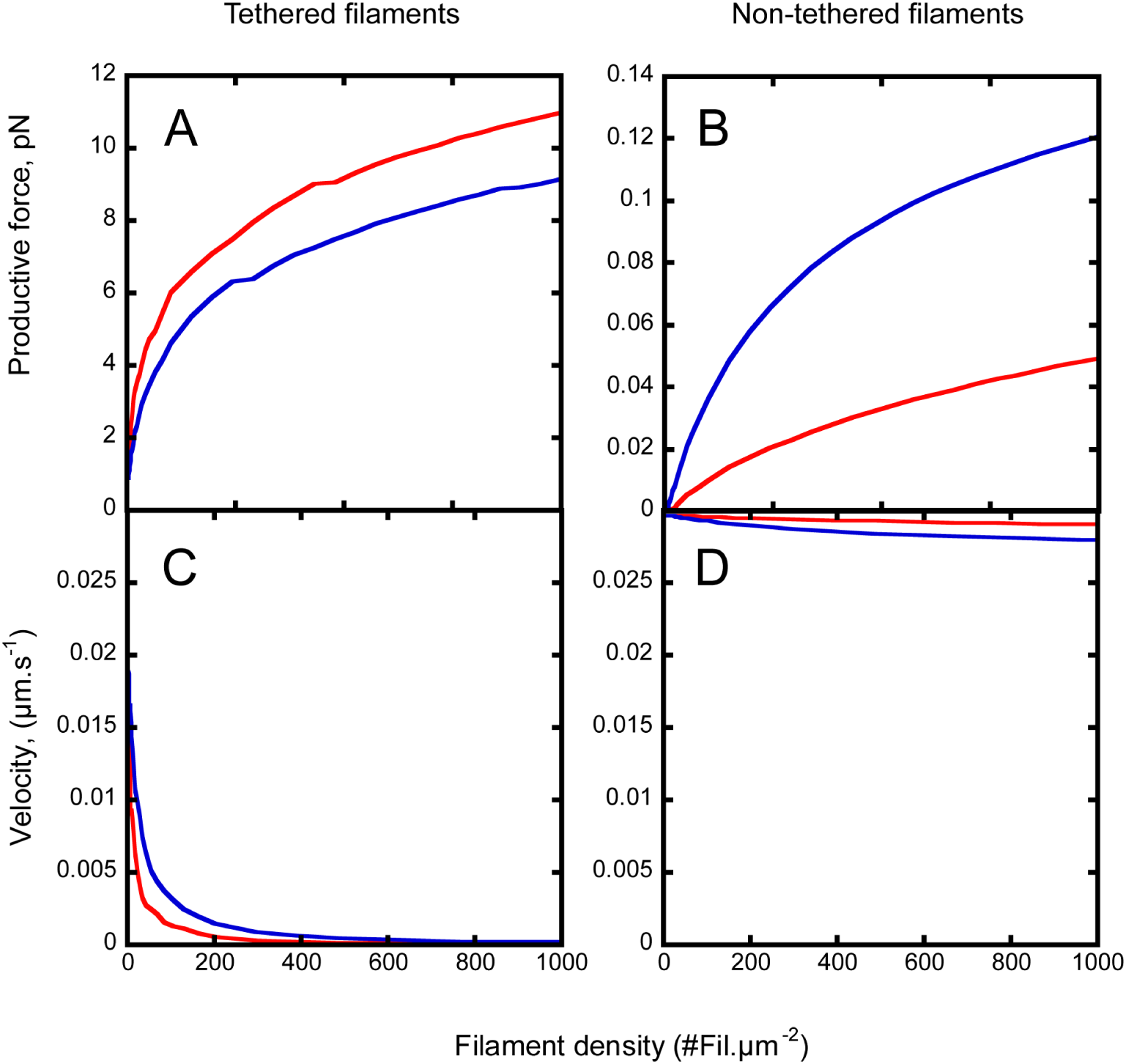
Force developed by a bundle of filaments. <u>Panels A and B:</u> 2-µm long tethered (A) or non-tethered (B) filaments exert a force which scales with filament density in the bundle. Note that attachment conditions yield a difference of ∼two orders of magnitude for the force exerted by the filaments. <u>Panels C and D:</u> The bead velocity rapidly decreases with the filament density for tethered conditions, whereas it remains virtually constant for non-tethered conditions. For all panels: red curve, θ_tilt_ = 0°, blue curve, θ_tilt_ = 35°.

Because a net force applied on the barbed end reduces the polymerization rate (Eq. 3), the large difference in the force produced by tethered and non-tethered filaments has dramatic consequences for the velocity of the load (Figs. 7C and D). Indeed, tethered filaments have a reduced polymerization rate and move the load at lower velocity. At high filament density, the velocity represents only a fraction of the maximal possible speed (Fig. 7C). Conversely, non-tethered barbed ends experience very low force, and their elongation is almost unaltered by the mechanical constraints. Non-tethered filaments are constrained to push the load even though they may slide on the load (Fig. 5B). In consequence, the load velocity is close to the maximal limit (Fig. 8D).

**Figure 8.**
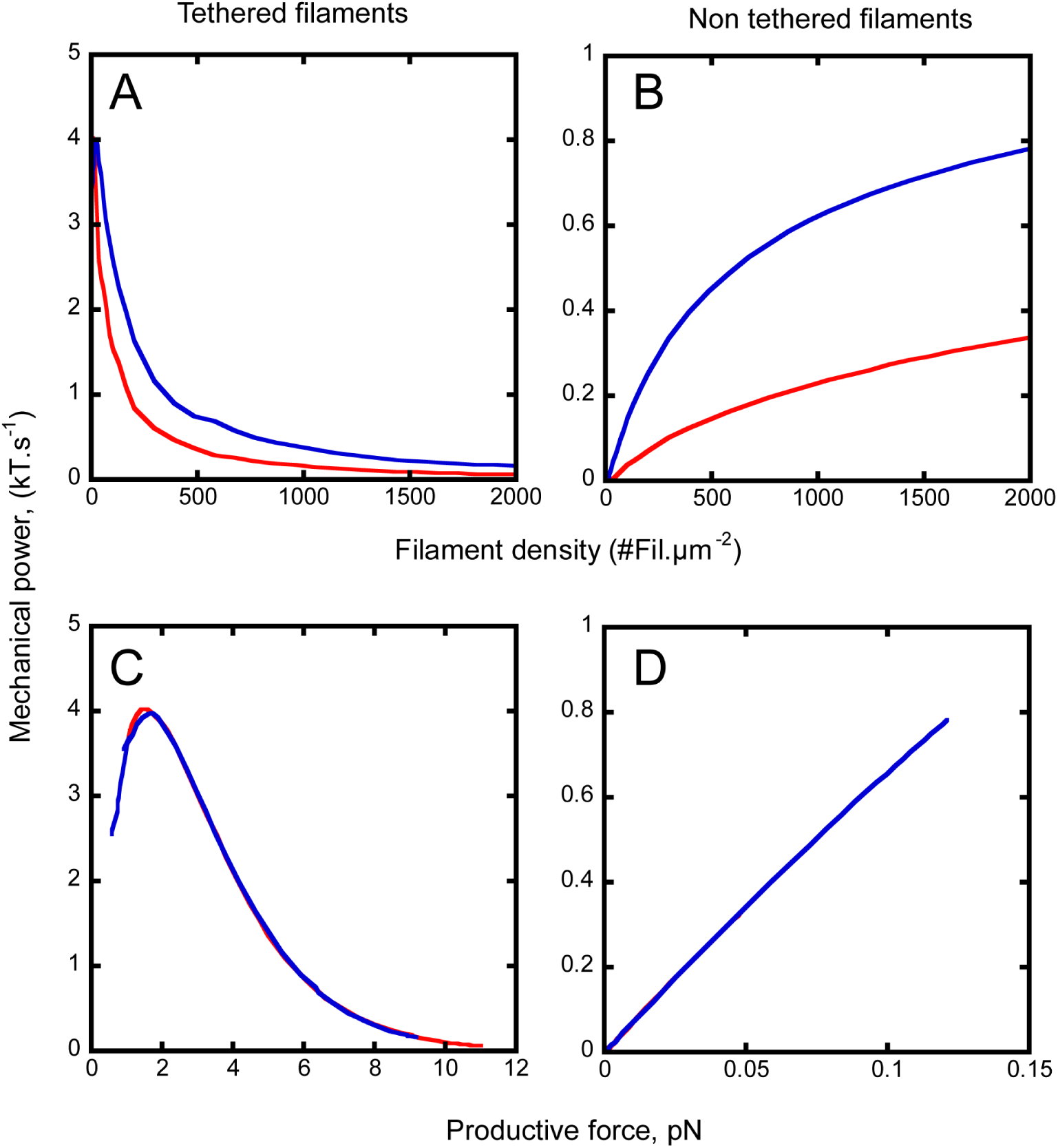
Mechanical power developed by a bundle of actin filaments. <u>Panels A and B:</u> Mechanical power transferred from the actin polymerization machinery to the load for tethered filaments (A) or non-tethered filaments (B) versus filament density in the bundle. <u>Panels C and</u> D: Mechanical power in function of the force produced by tethered (C) or non-tethered filaments (D). For all panels, simulation uses Eqs. 1-2 and 20-29 with 2-µm long filaments and persistence length of 10 µm; the free monomer concentration is 1 µM; red curve, θ_tilt_ = 0°, blue curve, θ_tilt_ = 35°.

### Mechanical power developed by actin filaments embedded in large filament populations

It might seem paradoxical that non-tethered filaments produce forces 2 orders of magnitude smaller than tethered filaments but move beads one order of magnitude faster (Fig.7). The mechanical power delivered by a single filament to the load, i.e. the product of velocity by force, is a more relevant metric to characterize the capacity of filaments to convert chemical energy into mechanical work (Fig. 8). The magnitude of the power and its dependence on filament density and force are dramatically different for tethered and non-tethered filaments. Non-tethered filaments have a power at least one order of magnitude smaller than tethered ones (∼0.8 kT.s^−1^ vs. ∼4 kT.s^−1^, Fig. 8). In addition, tethered filaments exert maximum power when the filament density is low (from 0 to 500 filament. µm^−2^) and is virtually independent from filament orientation (Figs. 8A and C), whereas the power of non-tethered filaments increases with both filament density and filament orientation (Figs. 8B and D). Strikingly, the force-power relationship for tethered filaments presents an optimal regime when filaments exert force around 1.5 pN and supply 4 kT J.s^−1^ to the load. Increasing the tilt angle does not change the maximal power and the force level at which this maximum is reached (4 kT.s^−1^ and ∼2 pN, Fig. 7 left bottom panel). In sharp contrast, non-tethered filaments do not present an optimal regime and their force-power relationship increases linearly with filament densities (Figs. 8B and D), and, surprisingly, does not depend on the orientation of filament pointed ends. In this respect, the force-power relationship of filaments in a bundle present universal features that depend on the attachment conditions at the barbed end only (Figs. 8C and D).

### Filament density and barbed-end attachment conditions control the bundle size

For non-tethered filaments, the physical interactions between the filament and the load remain unchanged, whatever the magnitude of the longitudinal force. Conversely, tethered condition requires tangent forces necessary to constrain the barbed end at a fixed position (Fig. 9A), and the tangent force magnitude *F_T_* varies moderately with the tilt angle (Fig.9, red and blue curves). If we model the tether as a spring with stiffness parameter *k_stiff_* and zero equilibrium length, its extension under load is given by *X_Link_* = *F_T_*/*k_stif f_* (Fig. 9B). When the tether is in a thermal bath, the probability for such an extended link to exist is (see Fig. 10C)

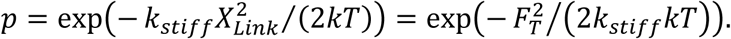

**Figure 9.**
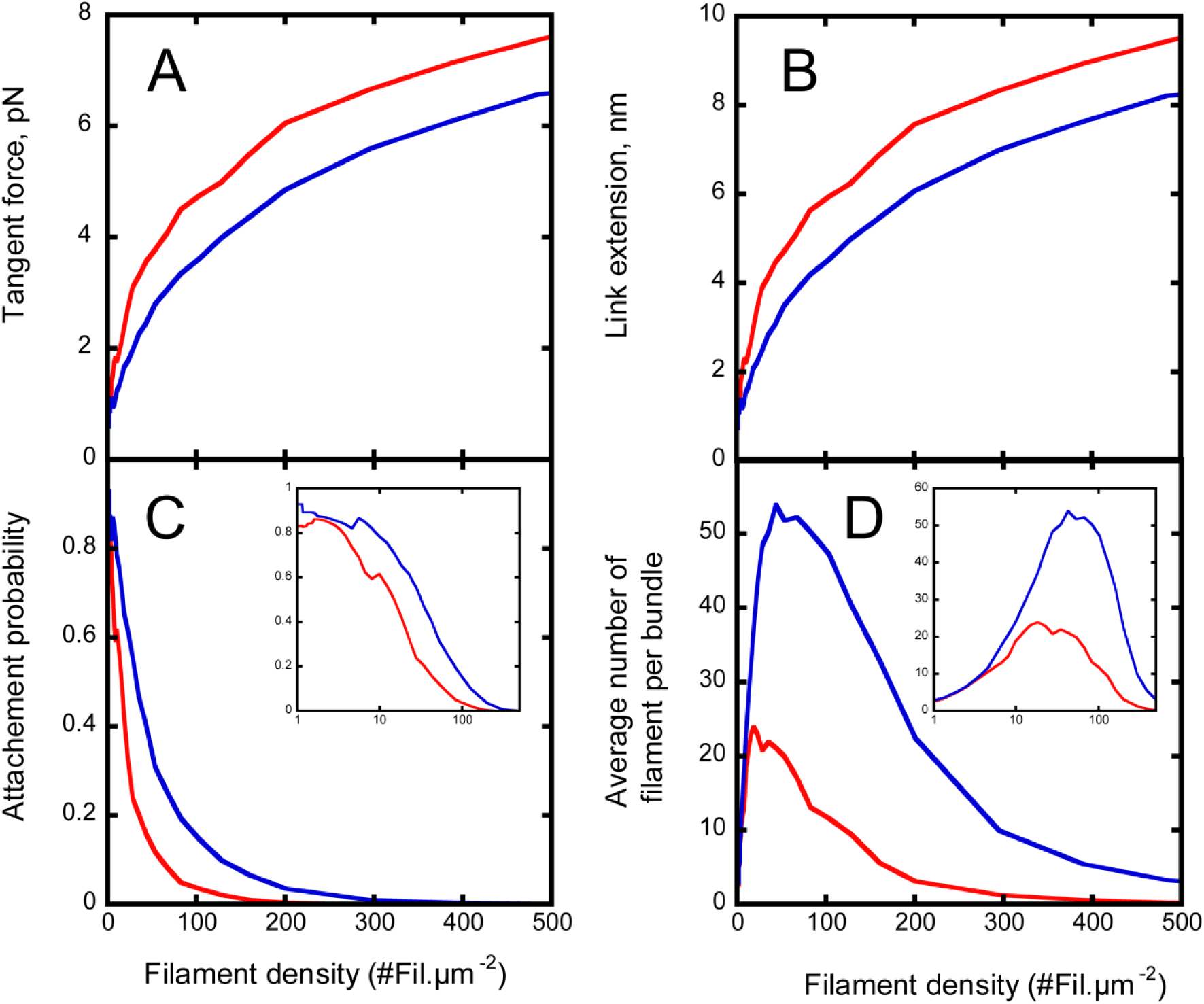
Average number of filaments in bundles when barbed ends are tethered. <u>Panel A:</u> Tangent force exterted on the barbed end of actin filaments for 0° tilt (red curve) and 35° tilt (blue curve). <u>Panel B:</u> Link extension at equilibrium for a link stiffness *k_sti f f_* = 4 × 10^−4^*N. m*^−1^. <u>Panel C:</u> Probability for a given link to remain attached. <u>Panel D:</u> Average number of filaments per bundle. (C) and (D) insets: log-scale representation of the probability (C) and the average number of filaments per bundle (D) for lower filament densities.

By combining this probability with the density of filaments, we computed the average size of a bundle for a test attachment area of 1 µm^2^ (Fig. 9D). Our data show that bundle of tethered filaments can exist at low filament density only, and coincides with the densities where the transformation of chemical energy into mechanical power is maximal (Fig. 8). In these conditions, the average number of filaments per bundles remains quite low (∼20 to 50) and compares well with values found in our experiments (Fig. 2). For higher filament densities, the formation of stable bundles of tethered filaments is greatly inhibited by the very large constraints put on the tethers, which easily break them.

## Discussion

In this study, we have found simple, robust and universal mechanisms underlying the organization of bundles of actin filaments, their production of mechanical forces and their dynamical adaptation to external constraints. We demonstrated that lateral filament-filament interactions and the tethering of the filament to the load control the emergence of optimal conditions for force generation.

### Confinement, filament density and attachment of barbed ends control the force generated by filaments and creates the conditions for optimum mechanical power

We showed that the limitation of the amplitude of filament displacement favors wavy shapes and high magnitude force at the filament ends (Figs. 6, 7 and 8). However, this force increase upon confinement only happens for tethered filaments but not for untethered filaments. By preventing the filaments to slide on the load and to straighten their shape, the attachment constraint selects filament configurations that exert a high force onto their ends. Similar behavior has been demonstrated experimentally for intermediate filaments (35) and microtubules (36).

We also showed that confined bundles of filaments with tethered barbed ends have an optimal regime for the transformation of chemical energy into mechanical work (Fig. 8). This optimum, which is independent of the bundle orientation, occurs at moderate filament density where the lateral constraint is high enough to amplify the force exerted by the filament but sufficiently low to prevent a large reduction of the polymerization rate.

A simple calculation allows us to estimate the maximum efficiency of power production by filaments in a tethered bundle. In the ideal case where there is no viscous drag and no bending, the maximal filament elongation velocity is *V_max_* = *k_on_*[*G*]*δ*, and the polymerization force generated by the addition of a single subunit is *F* = (*kT*/*δ*)*log*([*G*]*/*[*G*]_0_). Hence, assuming a concentration of G-actin of 1 μM, the maximal power per filament is (Fig. 9C) *FV_max_* = (*kTk_on_*[*G*])*log*([*G*]/[*G*]_0_) ≈ 23 *kT*.s^−1^ (see Table 2), which implies that a single tethered filament works at 15% of the maximal yield only.

Our model also showed that tethered filaments are optimized to work at low filament density (Fig. S12, red curve) whereas non-tethered filaments develop their mechanical power when embedded in dense networks (Fig. S12, blue curve). Tethered and non-tethered conditions are mechanically equivalent at filament density of ∼ 400 Fil.µm^−2^. Therefore, tethered and non-tethered actin filament networks represent two optimal solutions to generate mechanical power over a broad spectrum of filament densities found in sub-cellular structures.

### Self-regulation of bundle size and organization

In physiological conditions, when a bundle of filaments is tethered to a load, the forces applied on each tether increases with the filament density in the bundle (Fig. 9), and tethers break if their deformations become too large. In consequence, the size of the bundle is self-regulated by the filament density in the network and is maximum at moderate network density (Figs. 9 and S12).

The prediction of an optimum for the mechanical yield and the bundle size (Fig. 9) are two independent outputs of our biophysical analysis. It is important to stress that both optima occur for tethered filaments embedded in networks of moderate density (100 to 500 filaments per µm^2^), in close agreement with densities present in cells estimated at 400 to 1000 filament per µm^2^, as measured in electron micrographs (37).

### Buckling as a mechanism to control filopodia length

Our results suggest that formin-driven extension of bundles, similar to the bundles in filopodia, is mainly controlled by bundle rigidity (Figs. 3 and 4), which is proportional to the number of active barbed ends. For example, when a filopodia reaches a critical length, which is determined by its filament density and attachment to the plasma membrane via formins and/or Ena/VASP, it stops pushing the membrane and buckles. This transition between extension and buckling (Fig. 4) yields a simple and robust way to control the extension of filopodia in cells, as seen *in vivo* (13, 38). This mechanism is also important for the creation and stability of cell-cell junction in cell tissues, particularly in the formation of villi of same size that ensure tissue coherence (13, 14).

### Buckling is a way to release elastic energy in networks and accompany cytoskeleton deformation

Both *in vitro* experiments and models have shown the importance of buckling in the contraction of disordered stress fibers (21) and in the final dismantling of the network (20). Our study suggests this principle may be extended to the whole cell itself. When a bundle of filaments is subjected to mechanical forces, from other cells, obstacles or external forces for example, its buckling under a critical load may constitute an initial response that could trigger a more complex signaling pathway in the cell. Then, differences in the bundle composition (e.g. by modulating the nature or amount of crosslinks, motor proteins, number of filaments), geometry (e.g. by modulation bundle length, anchoring at the plasma membrane or on an organelle) and/or mechanics would modulate the cell response and yield different fates for the cell cytoskeleton. When bundles are stiff (e.g. if they are crosslinked or composed of a large number of filaments), the elastic restitution of the energy stored in bundle buckling could allow cells to resist external constraints. For soft bundles (long ones, without crosslinkers or made of severed filaments), the stressed cytoskeleton would fall in rapidly. For example, it has been shown that spatial 3D distribution of bundles and their interactions (either bundle-bundle or bundle-rest of the cytoskeleton junctions) is crucial for cells (39). Last, bundles are part of the fiber system allowing cells to communicate with other cells (e.g. bundles in villi), to sense the extracellular space (e.g. filopodia) or to couple to focal adhesions. Therefore, any biological condition that change either the geometry or the mechanics of bundles could exert control over cell dynamics.

## Acknowledgements

JB was supported in part by the National Institutes of Health/National Institute of General Medical Sciences Grant R01GM115636. LB is supported by an ANR MaxForce (ANR-14-CE11- 0012-02).

## Supplementary material

### A. Simplified model for the analysis of the experiments

Correct interpretation of experimental observations in Fig. 1 requires the determination of actin filament bundle rigidity. We limit our model for bundle mechanics and deformations to 2D-bending strain. This simplification is dictated by the available experimental data (TIRF microcopy images, Fig. 2) which give access to 2D bundle deformations only. Therefore, we assume that bundle rigidity is given by a single parameter, the apparent rigidity modulus (or persistence length) which depends on two quantities: the bending rigidity of a single filament and the number of filaments in a bundle (1). We focus our analysis on bundles which self-interact to form a stem and a loop (Fig. 2). These configurations represent structures at mechanical equilibrium when elastic forces, which tend to straighten the bundle, are balanced by attractive (or depletion) forces that keep distant sections of the bundle to form a stem (Fig. 2).

#### a. 2D equations for the mechanical equilibrium of filaments lying in a plane

To determine the rigidity of filament bundles (Fig. 1A), we adapt Eqs. 1-2 to 2D actin filament bundles as observed in TIRF microcopy (Fig. S1B). The material frame vectors (**d**_1_, **d**_3_) are in the plane (**d**_2_ points out of the plane). The orientation of the material frame (**d**_1_, **d**_3_) requires a single parameter, *θ*, the angle between **d**_3_ and the horizontal axis. The filament strain or curvature d*θ*/*ds* ds enters in the definition of bending strain vector as 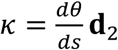. From Eq. 1, we derive the force and moment balance equation in its component-wise form

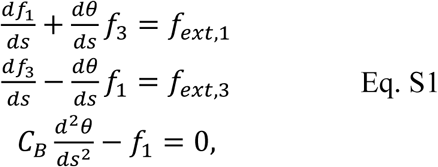

where *f_i_* is the component of the internal force along the director **d**_i_, i=1, 3, and *C_B_* is the bending rigidity. Note that the bending rigidity parameter, *C*_B_, and the persistence length, *L*_p_, are related by C*_B_* = *kTL_p_* where *k* is the Boltzmann constant and *T* the temperature in Kelvin. The twist strain, which is proportional to *C_T_* in Eq. 1, is absent from Eq. S1. This approximation is valid since twist energy is always lower than bending energy (2) and that 2D filament mechanics is controlled by filament bending curvature (3). The inextensibility condition (Eq. 2) gives two equations for the horizontal and vertical components of **r***(s)*:

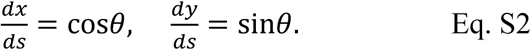

For filaments or bundles forming a loop (Fig. S2), the right hand side (*rhs)* of Eq. S1 (two first lines) models the filament-filament interactions responsible for loop formation and its stabilization (Figs. S2A and S2B). Because a loop is symmetric, we assume that the force between two points on the filament, **M** and **M**’, is directed along the line **MM’** and proportional to |**MM**| = |*x*| (Fig. S2C). The force is attractive as long the points are within a distance *r*_1_ from each other and repulsive if the distance is smaller than *r*_0_ (hard core repulsion). The force vanishes for inter-filament distance *x* larger than *r*_1_. Using these assumptions, Eqs. S1a and S1b are changed into:

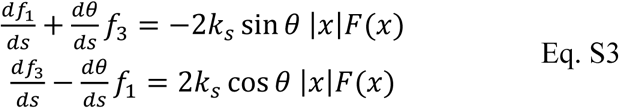

with

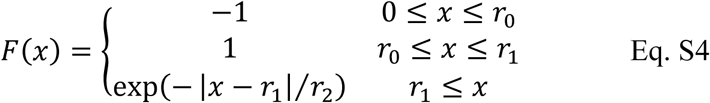

After normalization of the position (*x, y*) and arc length variable *s* by the filament length *L*, the final equations read

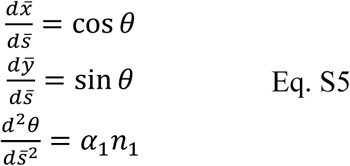

and

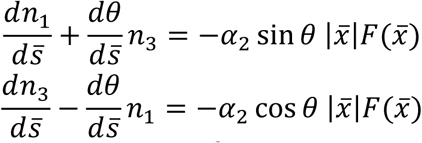

where 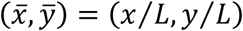, 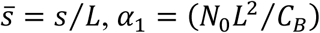, *α*_1_ = (*N*_0_*L*^2^/*C_B_*) and *α*_2_ = (2 *k_s_L*^2^/*N*_0_), *L* is the total loop contour length, *N*_0_ = (*kT/L*_0_) is the natural force unit for the system ( ≈ 4.1 × 10^−15^ *N*) with *L_0_*=1 µm, *n*_1_ and *n*_2_ are the force components normalized by *N*_0_. Note that α_1_ and α_2_ are dimensionless parameters. All the variables and parameters are summarized in Table S1. The vertical symmetry of the configuration (Fig. S2C) gives additional relationships

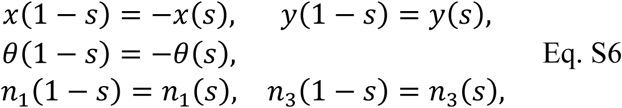

All these conditions are a direct consequence of the loop symmetry with respect to the vertical *y*-axis.

**Table S1.** Variables and parameters used in the analysis of bundle rigidity.

| Variables | Definition | Dimension | Typical value |
| --- | --- | --- | --- |
| $s$ | Arc-length | L | 20 to 100 $\mu\text{m}$ |
| $(x,y)$ | Point coordinates on the filament | L | |
| $\theta$ | Angle between the tangent and the horizontal axis | | |
| $(f_1, f_3)$ | Force along $\mathbf{d}_1$ and $\mathbf{d}_3$. | M.L.T<sup>-2</sup> | |
| **Parameters** |  |  |  |
| $kT$ | Thermal energy | M.L<sup>2</sup>.T<sup>-2</sup> | $4.1 \times 10^{-21}$ J |
| $k_s$ | Self-interaction force density | M.T<sup>-2</sup> | $2.7 \times 10^{-3}$ pN.$\mu\text{m}^{-2}$ |
| $r_1$ | cut-off distance | L | 0.055 $\mu\text{m}$ |
| $r_2$ | spatial decay for the self-interaction force. | L | 0.02 $\mu\text{m}$ |

#### b. Boundary conditions

Solutions of Eqs. S5 depends on the boundary conditions at the ends of the filament, i.e., at 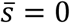 or 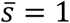. Since the attraction force exerted along the stem balances the elastic force due to the loop rigidity (Figs. S2A, S2B and S2C), the components of the internal force (*n*_1_, *n*_2_) should vanish at 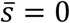

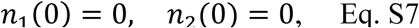

In consequence, from Eq. S6, the internal force should also vanishes at 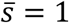. We complete the set of boundary conditions by specifying the position of the filament at *s*=0 and *s*=1

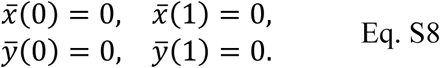

#### c. Determination of bundle bending rigidity

The shape of a loop is given by a solution of Eqs. S5-S8 valid for a 2D-elastic rod. However, since solutions of Eq. S5 depend on the ratio (*α*_1_/*α*_2_) only, we cannot have access to bundle rigidity directly. Therefore, we first determined the force interaction in the case of a single filament for which the bending rigidity is known. Then, we used the same method to analyze the elasticity of loops formed by a bundle made of several filaments by scaling the parameters (*α*_1_, *α*_2_) appropriately.

##### Algorithm for the determination of the interaction parameter in the case of single filament loop (see Table S1)

1. We extracted the configuration of the loop and the stem from the microscopy image (Fig. S2A and S2B) and measured its contour length (*L*).
2. We normalized the filament configuration and arc-length with the filament contour length *L*.
3. Using a persistence length of 10 µm for a single actin filament (or a bending modulus *C_B_* = 4.1 × 10^−26^ *N. m*) (4, 5), and *L*, we estimated the parameter
4. *α*_1_ = (*N*_0_*L*^2^/*C_B_*) (with *N*_0_ = 4.1 × 10^−15^N)
5. The best configuration fit (Fig. S2D and S2E) was obtained by adjusting α_2_, the unique free parameter remaining in Eq. S5.
6. *k_s_* was then given by

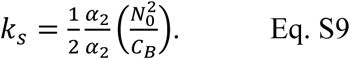

The self-interacting force density *k_S_* is attractive along the stem up to the junction between loop and stem sections (red arrows, Fig. S3). It peaks (∼100 pN.µm^−1^) at the transition between stem and loop where elastic forces from the loop counterpoise attraction in the stem (Fig. S3).

##### Determination of bundle rigidity

We now focus on the determination of the rigidity of a bundle made of several filaments. Firstly, assuming a close packing of filaments, the radius *R* of a bundle made of *N* filaments scales as 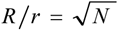, where *r* is the radius of a single filament (Fig. S4). Secondly, we assume that the interaction force is proportional to the perimeter of the bundle, since we expect that only the filaments in the outer part of the bundle can exert force on filaments outside the bundle (Fig. S4). Therefore, the parameter *k_S_* in Eq. S3 (or α_2_ in Eq. S5) scales as

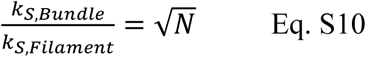

In addition, previous studies showed that simple geometrical arguments allow to approximate the apparent bundle rigidity as a power function of the number of filaments

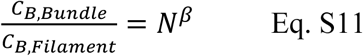

with 1≤ *β* ≤ 2 (1). The case *β=1* corresponds to filaments free to slide in the bundle, and *β=2* corresponds to totally crosslinked filaments. By combining Eqs. S10 and S11, we conclude that

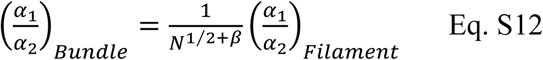

Therefore, by comparing experimental images of bundles forming a loop obtained by TIRF microscopy to a solution of Eqs. S5-S12, we can deduce *N*, the number of filament in a bundle, and, thanks to Eq. S11, the bundle rigidity. Fig. S6 shows typical observed closed loops (dotted lines) and the corresponding optimal solution to Eqs. S3-S6 and the parameters listed in Table S1.

#### d. Polymerization kinetics and polymerization force

In presence of profilin, the polymerization rate for actin filaments barbed ends capped by the processive formin mDia1 is 38 µM^−1^s^−1^ (6). Assuming a depolymerization rate of 0.1 s^−1^ (7), we predict the critical concentration for actin monomer to be [G]_0_ = 0.037 µM (Table S2). The concentration of actin monomers, which yields a constant elongation of 0.2 µm.min^−1^ for actin filaments (measured from Fig. 1 and Fig. S5), is [G]= 0.068 µM (Table S2). In consequence, the maximal force developed by actin polymerization, given by *F_pol_*_,*max*_ = (*kT/δ*)ln([*G*]/[*G*]_0_), is 0.93 pN (defined in Table S2).

##### Drag force

The presence of actin filaments in the bulk increases the apparent viscosity in the experiments close to values characteristic of that of a cell. Direct viscosity measurement *in vivo* reported values ranging from 0.08 to 0.26 Pa.s (8, 9) which represents a hundred-fold increase of viscosity compared to that of water (0.001 Pa.s). Given the polymerization force (0.93 pN) and drag force exerted on a sphere (Table S2), we predict that the drag force during bead movement is in the range 0.01 to 0.032 pN (Table S2).

##### Buckling force

The critical force required to buckle an elastic bundle is:

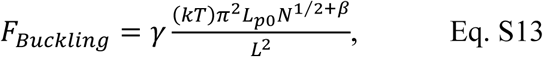

where *L_P_* is the persistence length of a single actin filament, *N* is the number of filaments in the bundle, *L* is the bundle length, and γ is a numerical factor that depends on the boundary conditions at the bundle ends [7]. In consequence, when the drag force balances the critical force at buckling

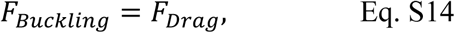

and the bead is stalled while bundle elongation continues. This transition occurs at a critical bundle length given by solving Eq. S14 with the help of Eq. S13

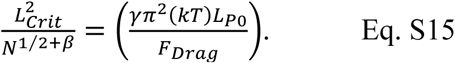

Note that the *rhs* of Eq. S15 is a constant which does not depend on bundle characteristics. The estimated values (Table S2) yield critical lengths in the range of experimental observations. From Eq. S15 and assuming constant elongation, one can predict that large bundles will sustain the bead propulsion regime for longer time periods.

**Table S2.** Parameters used to model bead motility

| Parameter | Definition | Value | Reference |
| --- | --- | --- | --- |
| $r$ | Bead radius | 2 $\mu\text{m}$ | |
| $\delta$ | Actin monomer radius | 2.75 nm | |
| $C_{B0}$ | Single actin filament bending rigidity | $4.1 \times 10^{-26} \text{ N.m}$ | |
| $V_{\text{max}}$ | Constant bead velocity | $0.0033 \mu\text{m.s}^{-1}$ | |
| $k_{\text{on}}$ | Polymerization rate (barbed end) in the presence of formin | $38 \mu\text{M}^{-1}.\text{s}^{-1}$ | (6) |
| $k_{\text{off}}$ | Dissociation rate from the barbed end | $1.4 \text{ s}^{-1}$ | (10) |
| $C$ | Drag coefficient, $C = 6\pi\eta r$ | $1.13 \times 10^{-7} \text{ Pa.s.m}$ | |
| $[G]_0$ | Critical actin monomer concentration ($k_{\text{off}}/k_{\text{on}}$) | $0.037 \mu\text{M}$ | |
| $[G]$ | Apparent monomer concentration | $0.067 \mu\text{M}$ | |
| | $[G] = [G]_0 + \frac{V_{\text{max}}}{k_{\text{on}}\delta}$ | | |
| $F_{\text{Pol}}$ | Polymerization force, | 0.93 pN | |
| | $F_{\text{pol,max}} = (kT/\delta)\ln([G]/[G]_0)$ | | |
| Mechanical power, $W$ | $W = F_{\text{pol}}V_{\text{max}} = (kTk_{\text{on}})[G]\ln([G]/[G]_0)$ | | |
| $\eta$ | Viscosity | 0.08 to 0.26 Pa.s | |
| Estimated drag force | $F_{\text{drag}} = (6\pi\eta r)V_{\text{max}}$ | 0.01 to 0.032 pN | |
| Critical buckling force (*) | $F_{\text{Buckling}} = \gamma \frac{(kT)\pi^2 L_{p0} N^{1/2+\beta}}{L^2}$ | unknown | |
| Critical length at buckling | $\frac{L_{\text{Crit}}^2}{N^{1/2+\beta}} = \left( \frac{\gamma\pi^2(kT)L_{p0}}{F_{\text{Drag}}} \right)$ | 40 to 12 $\mu\text{m}^2/\text{filament}^{1.5}$<br>(**) | |

(\*) $N$ is the number of actin filaments in the bundle; $L_{\text{Crit}}$ is the bundle length at the onset of buckling; $\gamma$ is a numerical factor.
(\*\*) using $\gamma=1$ and $\beta=1$ .

### B. Model of filament bundles during macroscopic bead movement

#### a. Equations used to simulate bead movements

Bead movement was modeled by a set of equations similar to Eq. S5-S8 without the external term accounting for loop formation

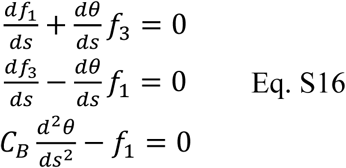

The arc-length variable, *s*, lies in the interval [0, *L*(t)] where *L*(t) is the time-dependent bundle length. Assuming that all filaments in the bundle experience the same stress at their barbed end, the global force-dependent elongation rate of the bundle is given by

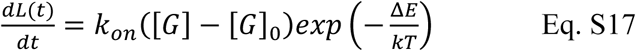

In the above equation, *k*on is the polymerization rate for actin filament capped by formins (6), [*G*] and [*G*]_0_ are, respectively, the concentration of actin monomers and the critical concentration in presence of formin; *δ* is the radius of a monomer, Δ*E* = *δ*|*f*_3_| is the work against the elastic force required to insert one monomer between the bead and the barbed end. The final set of equation governs the bead position

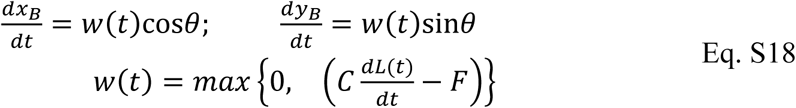

where (*x*_B_(t), *y*_B_(t)) is the bead center, *θ* is the angle between the bundle and the horizontal axis at the junction between the bundle and the bead, *F* is the magnitude of the constant viscous drag exerted on the bead, *C* is the coefficient giving the drag exerted on the bead (Table S2).

#### b. Boundary conditions used to simulate bead movements

The pointed end (at *s*=0) remains fixed in time with

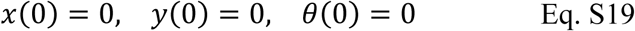

The boundary at *s=L*(t) connects the bundle and the bead

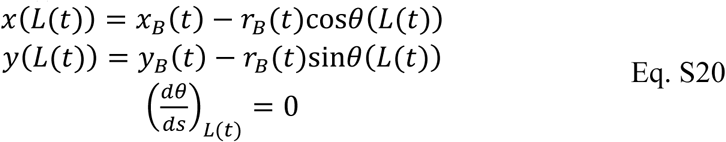

where *r*_B_ is the bead radius.

### C. Linearization of the 2D model

We start from Eqs. S1-S2 (Supplementary material, section A), which are valid for 2D systems (Fig. S1B). To simplify the analysis, we assume that

1. the angle *θ* between the tangent **d**_3_ and the horizontal axis is small;
2. the force applied along the tangent is zero (*f*_1_, _ext_=0)
3. the force applied laterally to the filament, which accounts for the presence of surrounding filaments, has a constant density (force per unit length), *f_Lat_.*

Eqs. S1-S2 read

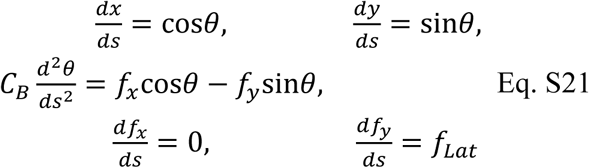

with (*f_x_*, *f_y_*) = (*f*_1_ cos *θ* + *f*_2_ sin *θ*,− *f*_1_ sin *θ* + *f*_2_ cos *θ*) and *C_B_* is the filament bending rigidity (*C_B_* = *kTL_p_*). After deriving the third line in Eq. S21 with respect to arc-length *s* and using the expression for the derivative of the forces (two last lines in Eq. S21), one gets

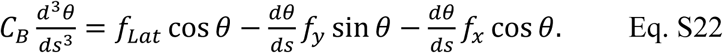

When θ is small enough, Eq. S22 is replaced by

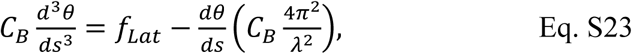

where we expressed *f_x_*, the constant unknown force along the horizontal axis, as

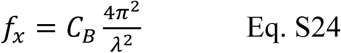

where *λ*, which has the dimension of the a length, is the *wavelength* of the wavy filament configuration (Fig. S8), the filament *wavenumber* ω (i.e. the number of bumps) is given by *ω* = 2*π*/*λ* (Fig. S8). The solution of Eq. S23 reads

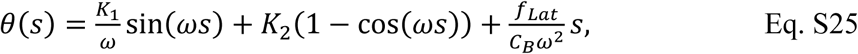

where we used the condition *θ*(0) = 0 to eliminate one of the arbitrary constants, *θ*(0) = 0 expresses that the initial filament direction is horizontal at the pointed end. The three unknown parameters (*K*_1_, *K*_2_, *ω*) are dependent on the conditions at the barbed end.

#### a. Tethered conditions

Since the filament is assumed straight and horizontal, tethered conditions are equivalent to

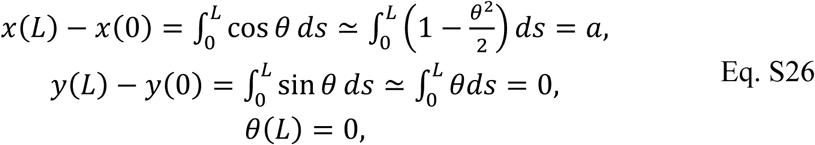

where *L* is the filament length and *a* the end-to-end distance (*a<L*). The first condition in Eq. S26 comes from the integration of the first line in Eq. S21 using the simplification cos *θ* ≃ 1−*θ*^2^/2, the second condition is obtained from the integration of the second line in Eq. S21 (with the approximation sin *θ* ≃ *θ*), while the third condition in Eq. S26 requires that the filament orientation at the barbed end be horizontal. Eqs. S25-S26 result in a set of non-linear equations for the unknown parameters (*K*_1_, *K*_2_, *ω*)

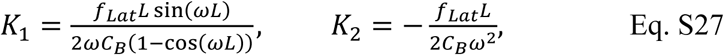

with ω solution of

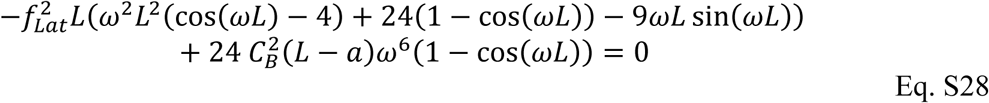

The integrated curvature is

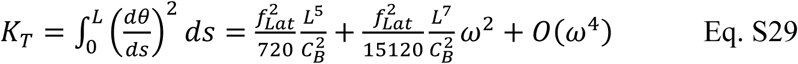

#### b. Non-tethered conditions

Non-tethered filaments have zero bending curvature at their both ends (Fig. 5B)

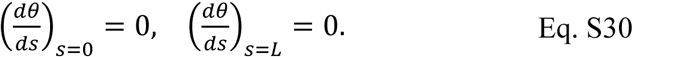

In addition, the tangential force *f_y_(s)* at *s=L* should be zero (Fig. 5B). The expression for *f_y_(s)* is derived from the third line in Eq. S21

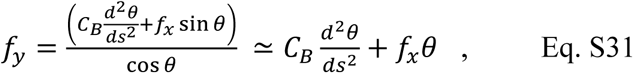

from which we get the last condition

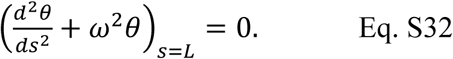

The solution of Eqs. S29 and S31 is

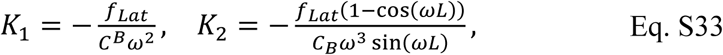

with ω given by the non-linear equation

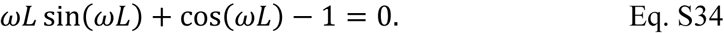

The associated total curvature reads:

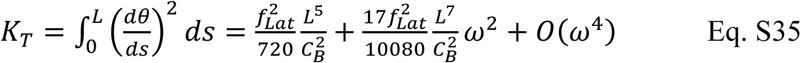

Also note that

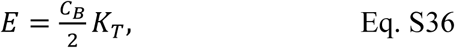

gives the total elastic energy stored in the filament shape for both attachment conditions.

#### c. Simulations

Once the lateral force density *f_Lat_* is fixed, the solution of Eqs. S27-S28 or Eqs. S23-S24 gives the wavenumbers ω and wavelengths *λ* = 2*π/ω* (Fig. S10). For a single value of *f_Lat_* there exists an infinite number of wavenumbers/wavelengths compatible with tethered (red curve) or non-tethered (green curves) conditions (Fig. S9). Which ω corresponds to the effective wavelength? For a given *f_Lat_*, the elastic energy stored in the filament shape scales with ω^2^ (Eqs. S29, S35, S36). Therefore, in realistic conditions, only the ground state, which corresponds to solutions with minimal ω (solution of minimal elastic energy), is observable. Filament configuration with higher ω cannot live in the noisy environment of the cytoplasm or *in vitro* experiments. From Fig. S9, we extracted the wavenumbers and wavelengths corresponding to the ground state and displayed them in Fig. 6.

## Supplemental figures

**Figure S1.**
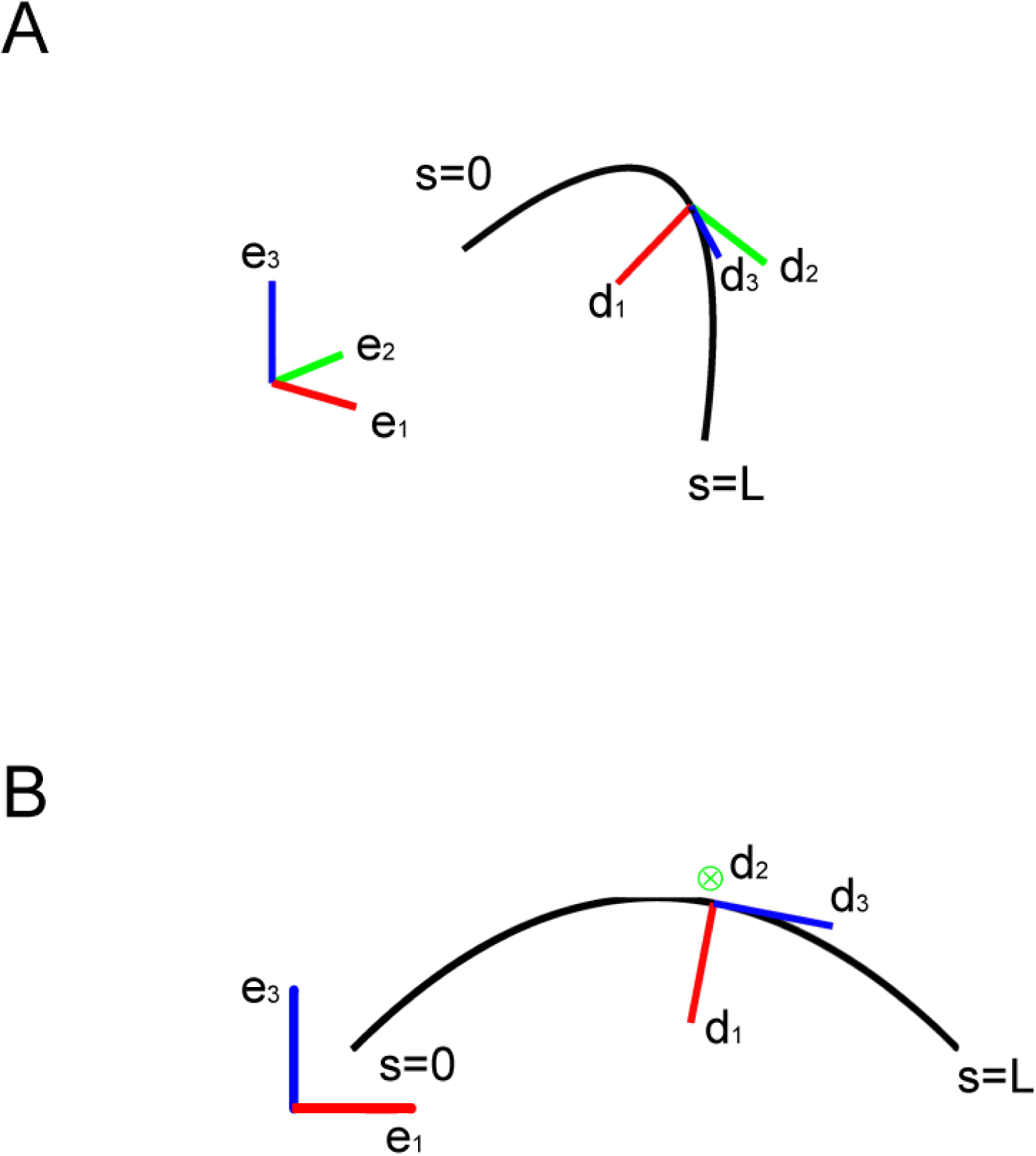
Schematic representation of 3D (A) and 2D (B) models for elastic filaments. (**e**_1_, **e**_2_, **e**_3_) (resp. (**e**_1_, **e**_3_)) is the spatial frame in 3D (resp. 2D representation). The directors vectors (**d**_1_, **d**_2_, **d**_3_) give the local orientation of the filament at any point. Note that in the 2D model, the bending moment is along **d**_2_ which is orthogonal to the plane (**e**_1_, **e**_3_).

**Figure S2.**
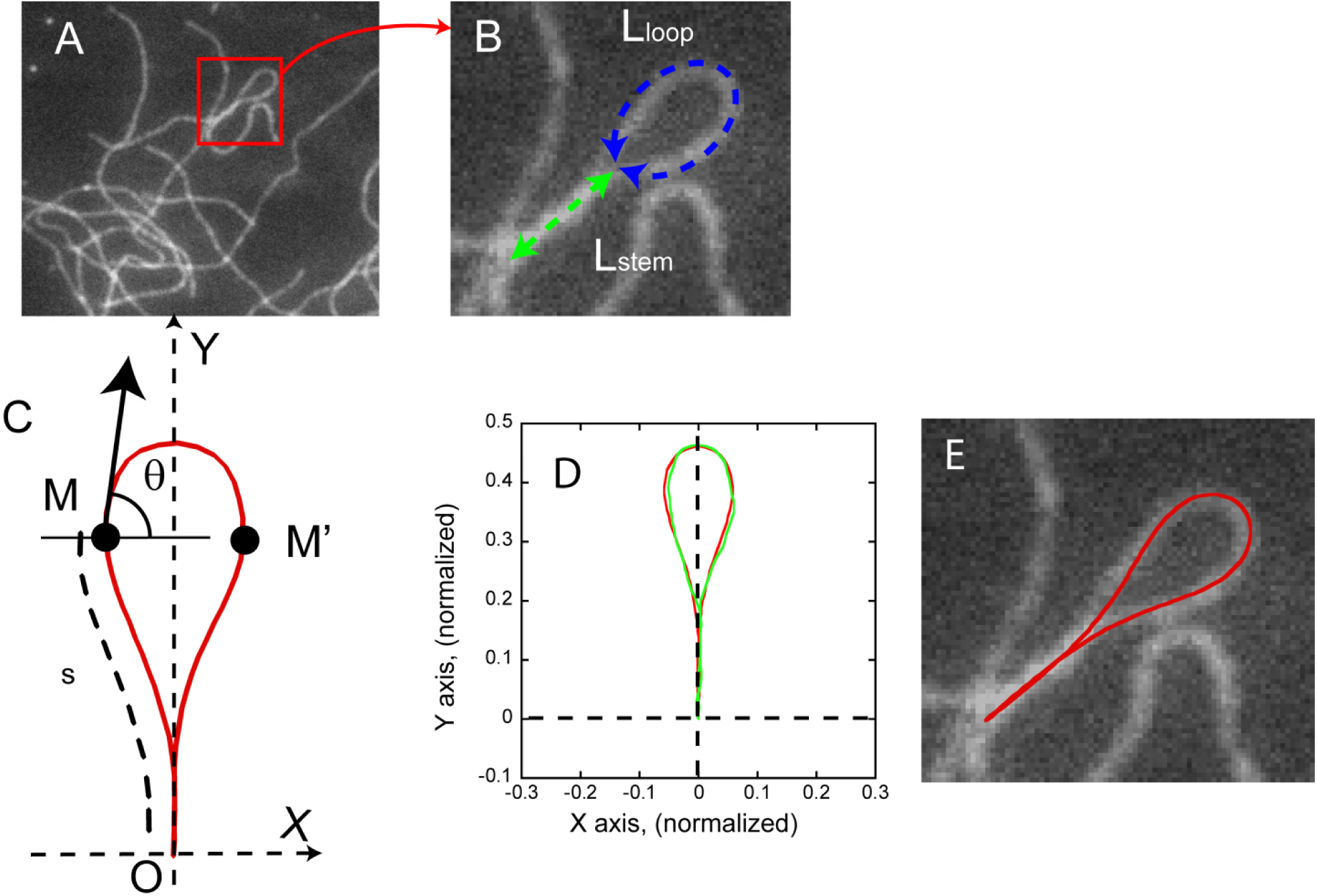
Determination of the attraction force: case of a single filament. <u>Panels A and B:</u> A folded filament forms a flat loop which is stabilized by attractive forces along the stem (dashed green arrow, panel B) balancing the elastic forces along the loop (dashed blue arrow, panel B). <u>Panel C:</u> Geometric model of the filament configuration in panel A. **M** is a point at distance *s* from the loop origin **O**, *θ* is the angle between the unit vector tangent to the filament at **M** (black arrow) and the horizontal axis. To simplify the model, we consider that the attraction force exerted on **M** is horizontal, along the line **MM**’. **M**’ is the point symmetric to **M** located at a distance *L-s* from **O**. Additionally, we assume that the attraction between **M** and **M**’ depends on |**MM**’| only. <u>Panel D:</u> Solution to Eqs. S3-S6 (red curve) superimposed to the actual filament configuration in A (green curve). Parameters of the solution are: *L*_f_ = 19 µm (total filament length); *L*_p_ = 10 µm (persistence length), k_s_ = 2.7×10^−3^ pN.µm^−2^ (self-interaction parameter), x_0_ = 0.055 µm and x_1_ = 0.05 µm. <u>Panel E:</u> Superposition of the solution shown in panel E (after rotation and scaling) and the filament.

**Figure S3.**
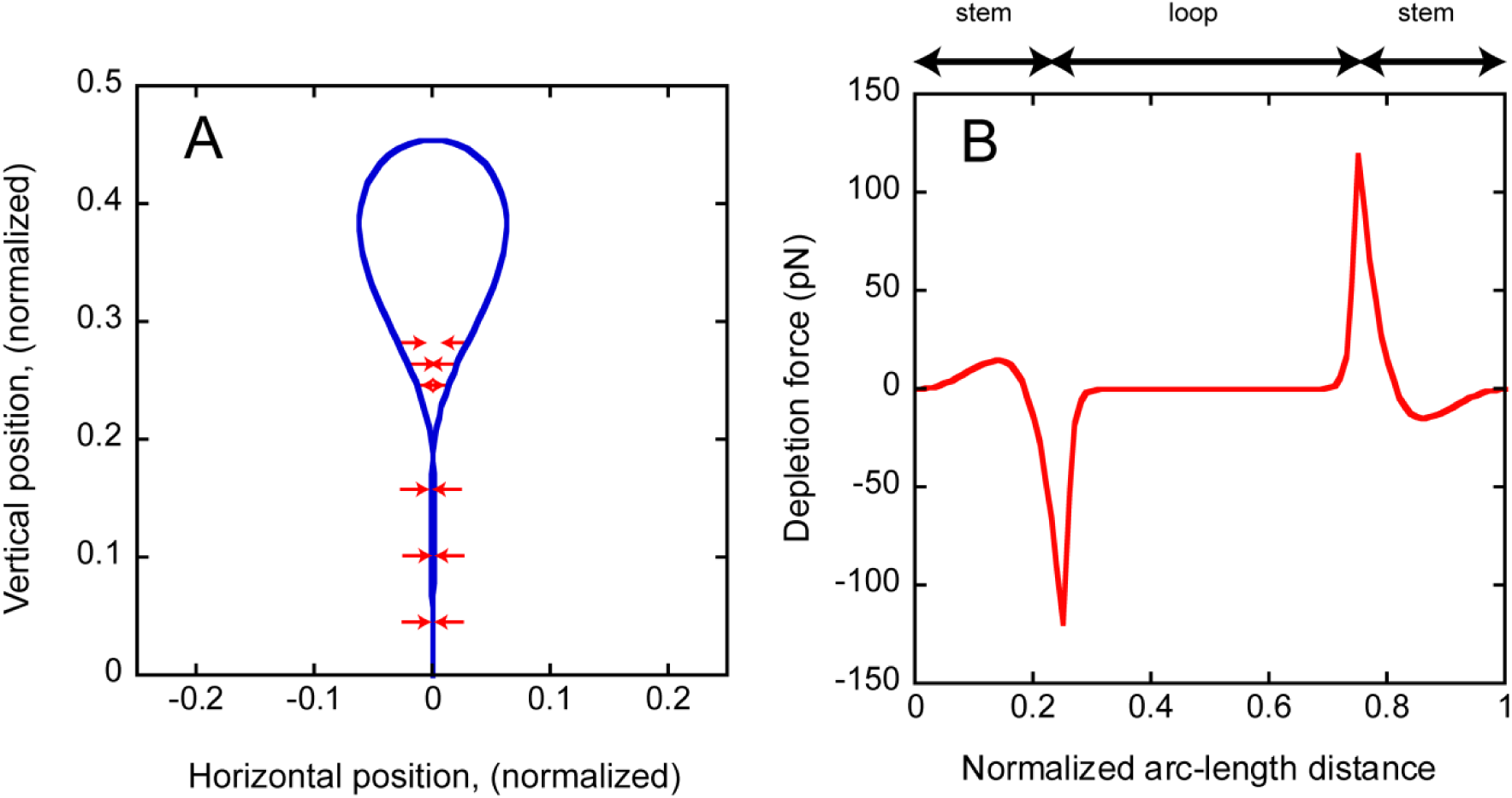
Attractive force density. <u>Panel A:</u> Schematic representation of the magnitude and sign of the attraction force. <u>Panel B:</u> Repulsion/attraction horizontal force as a function of the arc-length position for the solution shown in Fig. S2D.

**Figure S4.**
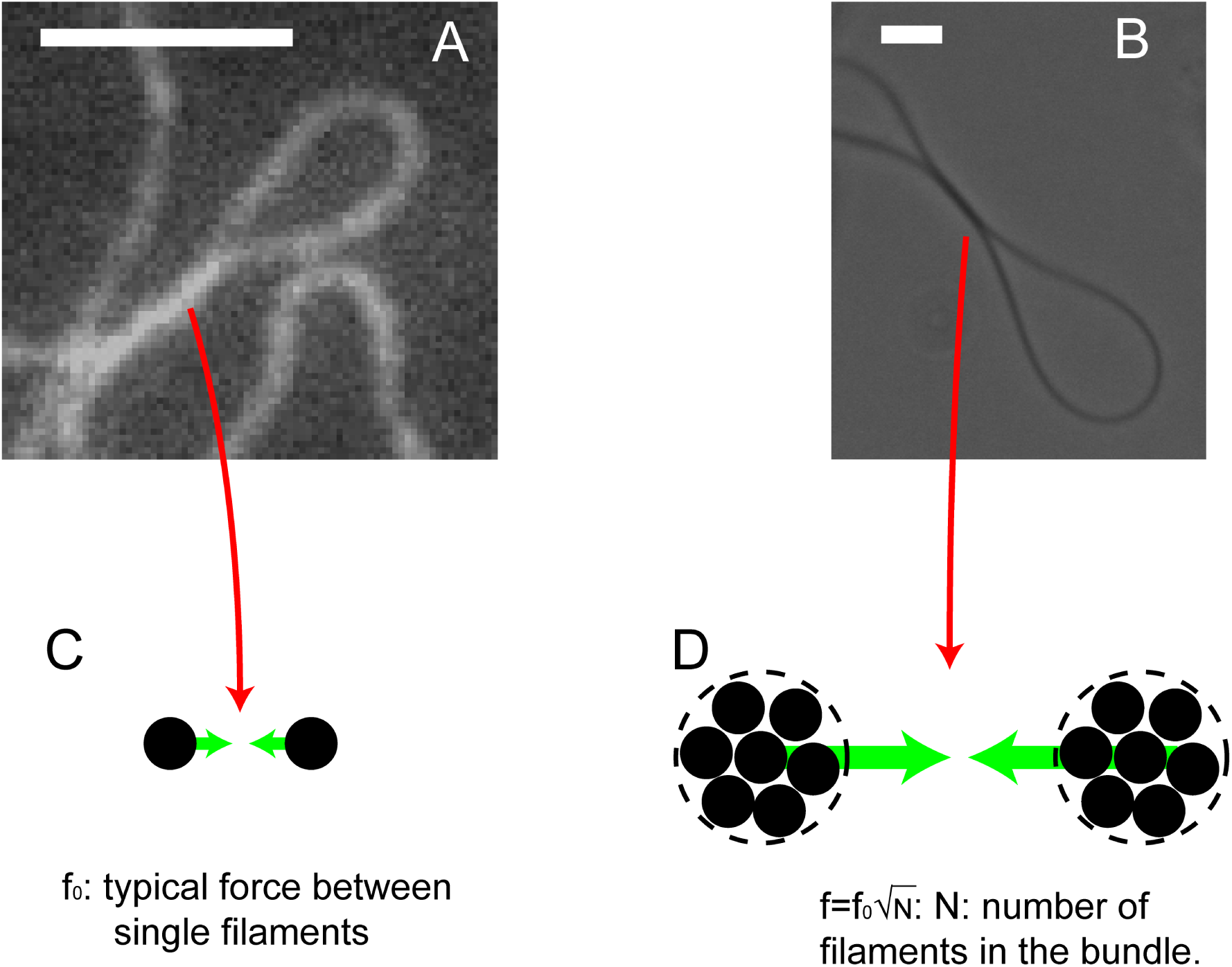
Model for the attractive force in bundles. <u>Panels A and B:</u> Typical loops made by a single filament (A) or a bundle (B). Scale bar: 5 µm. <u>Panels C and D:</u> Attraction force between filaments or bundles (green arrows). Small dark circles represent the cross section of a single actin filament (C) or actin filaments in the bundle (D).

**Figure S5.**
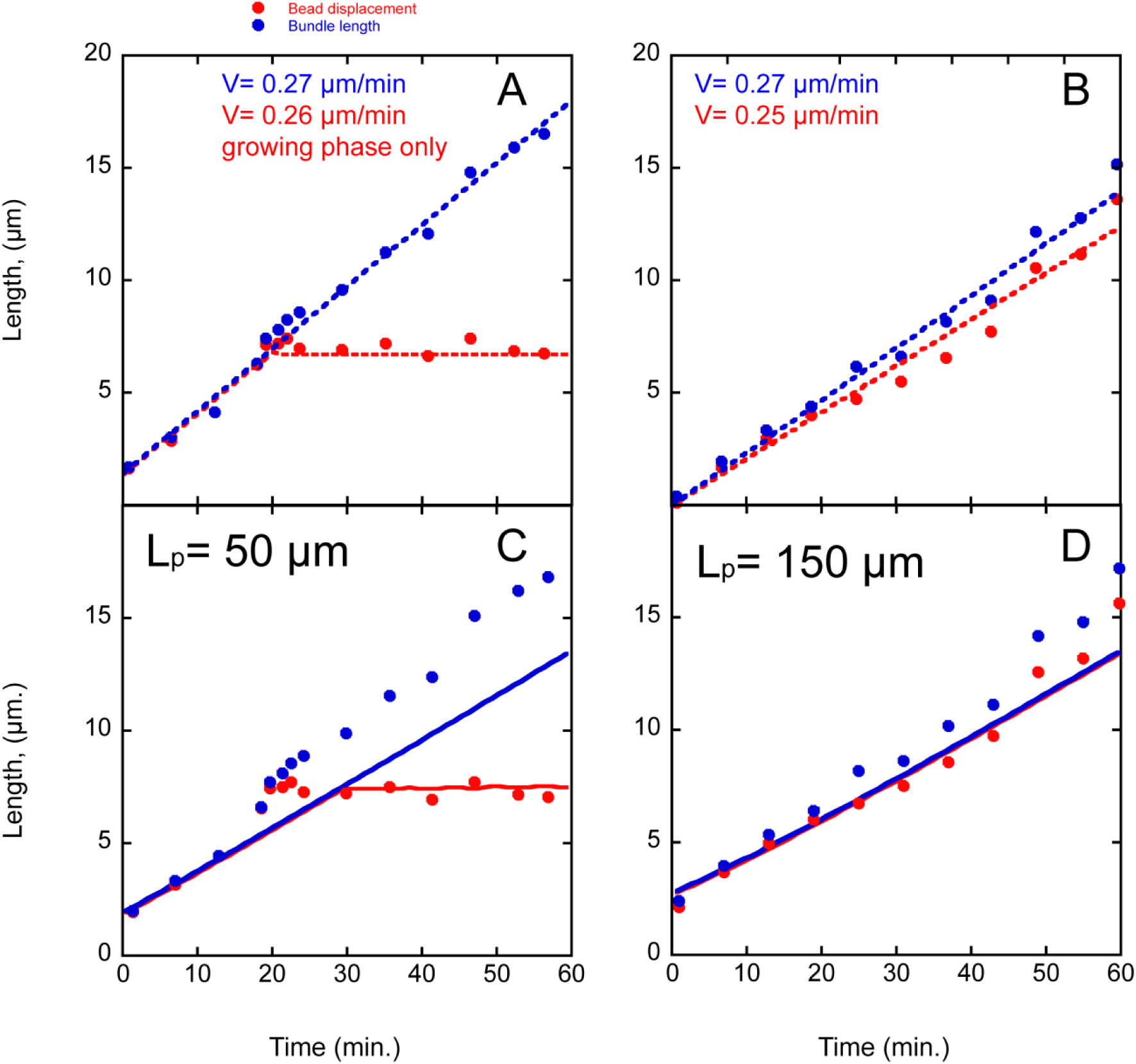
Kinetics of bundle elongation and bead displacement. <u>Panels A and B:</u> The bundle length (blue dots) and the bead trajectory (red dots) for the two are displayed for the two examples shown in Fig. 1A (red, bottom bead; blue, top bead). The dashed curves represent linear interpolation of experimental points. <u>Panels C and D:</u> Simulation of the bead displacement and bundle elongation using Eqs. S16-S20 for two bundle rigidities: (C) 50 µm, (D) 150 µm.

**Figure S6.**
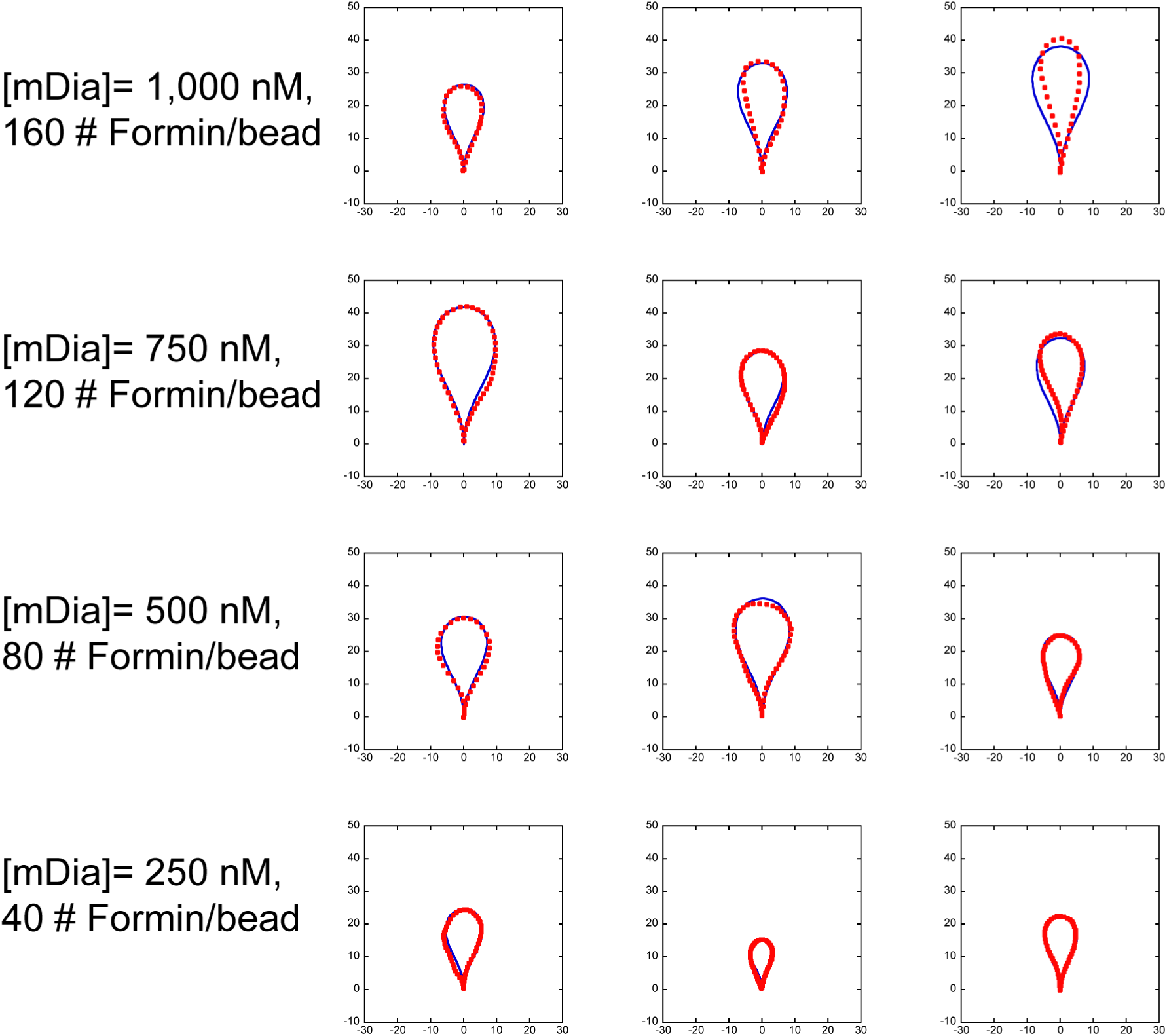
Sample of bundle loops for different density of formin. The blue curves give the best fit by Eqs. S5-S8 (with *β*=1) to the actual loop, shown in red. Horizontal and vertical axis are in micrometers.

**Figure S7.**
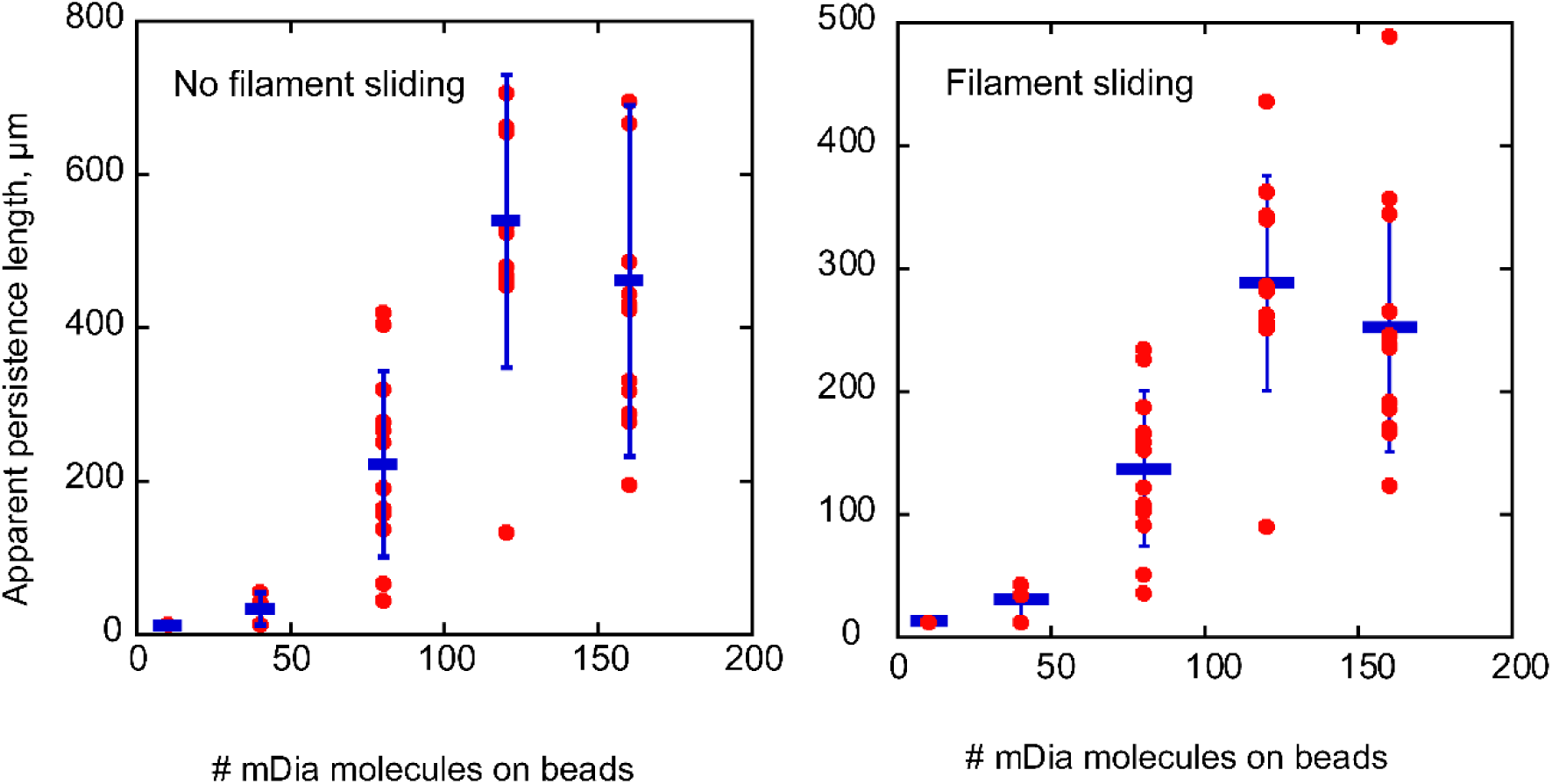
Persistence length of bundles. Determination of bundle persistence length using Eqs. S5-S8 assuming perfect sliding of the filament in the bundle (*β= 1*, right panel) or filament bound together (*β=2*, left panel). The average and standard deviation are indicated in blue.

**Figure S8.**
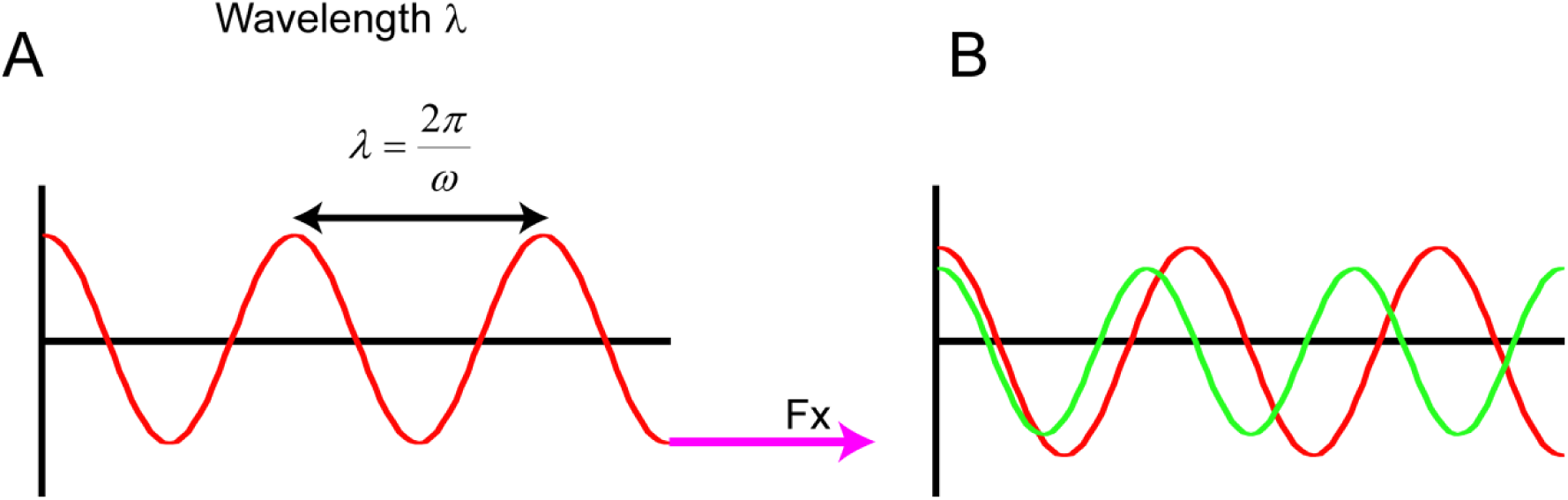
Wavy filament shape, wavenumber and wavelength. <u>Panel A:</u> The filament shape predicted by the model consists in a linear combination of periodic functions with period *λ* = 2*π*/*ω*. The force exerted by the filament onto its attachment point (magenta arrow) and the elastic energy stored in the filament shape are both proportional to ω^2^. <u>Panel B:</u> Because the number of half periods can change by steps of ±1 only, it results a sudden change in the wavenumber (henceforth in the energy and force).

**Figure S9.**
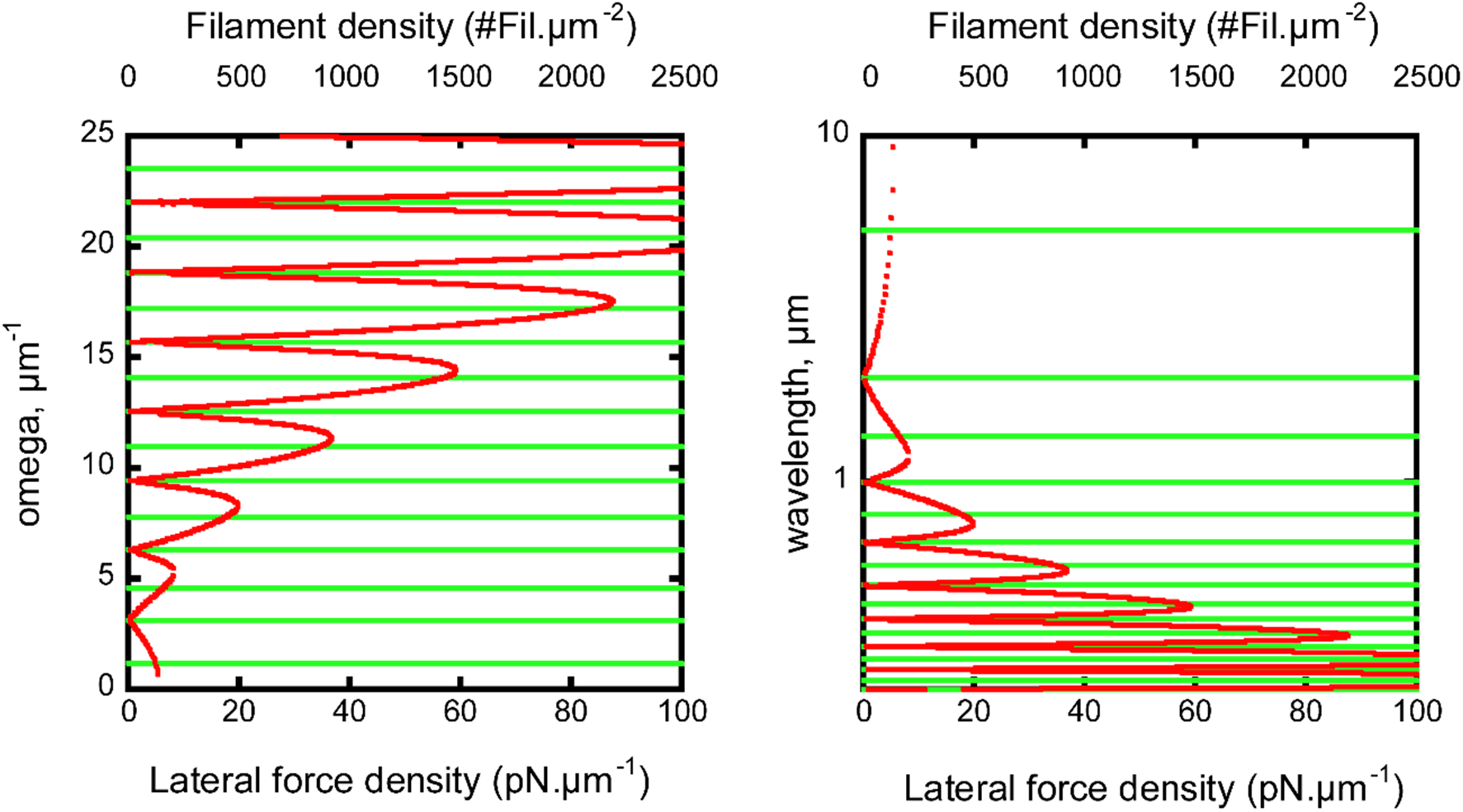
Filament wavenumber and wavelength predicted by the linear model. The wavenumber (left panel) and the wavelength (right panel) of filament configurations are computed as a function of the density of lateral forces coercing the filament into a cylinder. Red (resp. green) curves are obtained for tethered (resp. non-tethered) conditions.

**Figure S10.**
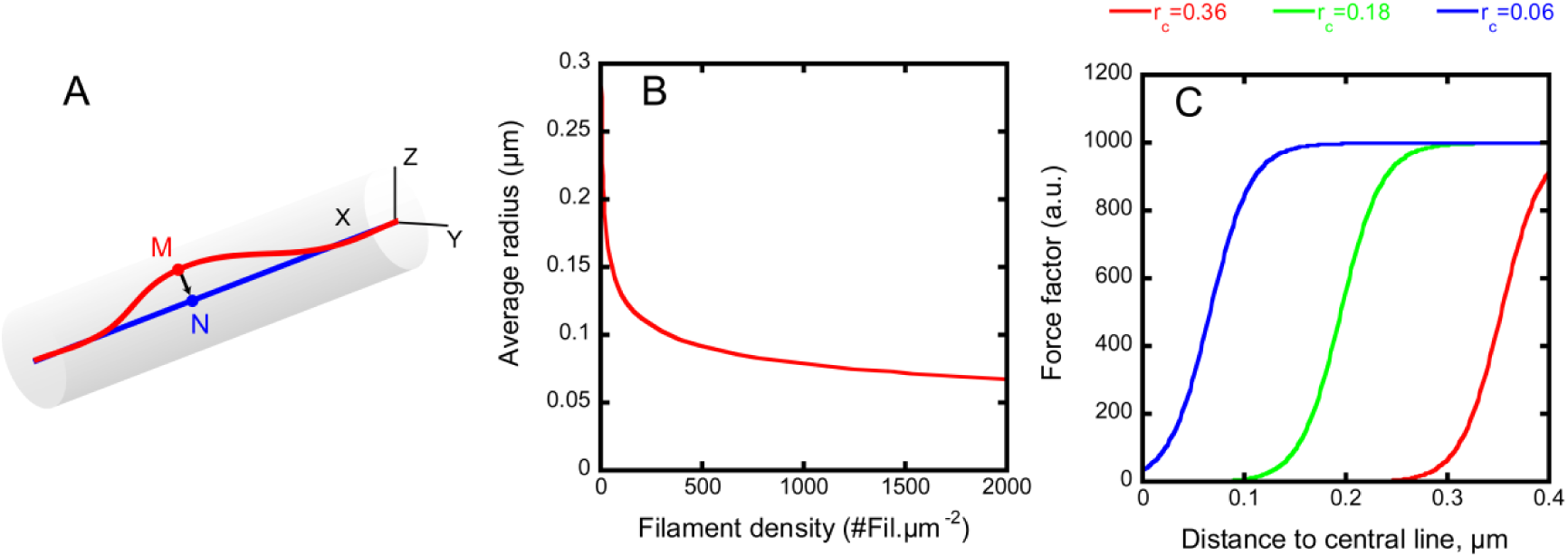
Model for filament in dense networks. <u>Panel A:</u> The force due to the presence of other filaments of the bundle is applied at any point M on the filament (red curve) along a direction orthogonal to the end-to-end axis (blue line). The gray cylindrical box represents the space in which the red filament is confined. The end-to-end blue line represents the filament configuration if the coercing force were infinite. <u>Panel B:</u> The radius of the cylindrical box containing the filament is a decreasing function of the filament density. <u>Panel C:</u> The amplitude of the force is given by the formula: *f* = (1 + exp((*r* − *r*_0_)/*r_c_*))^−1^ where *r* is the distance |MN| (see panel A) and *r_0_*, the box radius given in panel B. The three radii *r_0_*=0.36, 0.28 and 0.06 µm correspond to low (1 Fil.µm^−2^), moderate (50 Fil.µm^−2^) and high (2000 Fil.µm^−2^) filament densities. The scaling factor *r_c_* controls the transition width. We performed simulations with *r_c_*=0.02 µm. Values of *r_c_* in the range 0.01 to 0.05 give similar results (qualitatively and quantitatively), showing that it is the position of the transition (controlled by *r_0_*) that matters.

**Figure S11.**
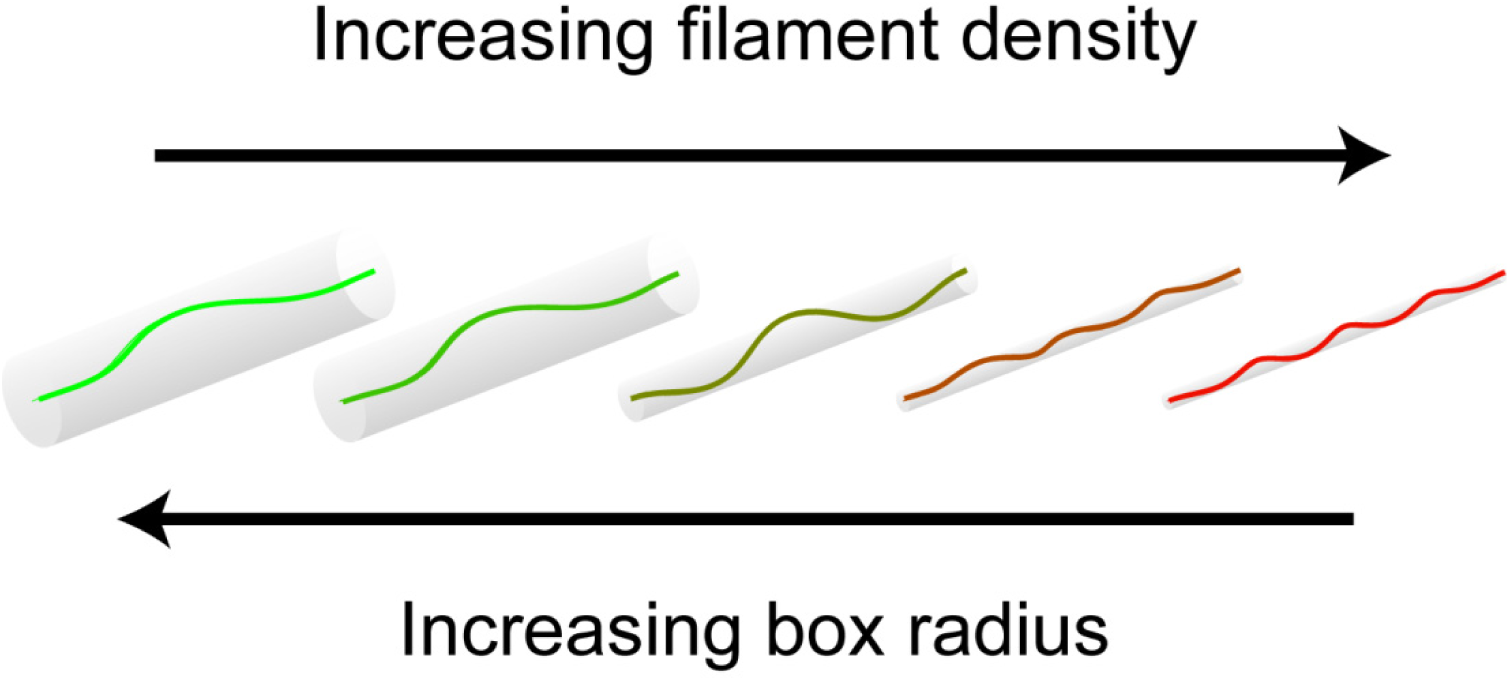
Actin filament is coerced into cylindrical box. Increasing the filament density from 1 to 2,000 filaments. µm^−2^ coerces a single filament into a cylindrical box with decreasing radius. The leftmost configuration (green curve) represents a buckled filament without lateral forces; the rightmost red configuration illustrates the presence of lateral constraints due to high filament density.

**Figure S12.**
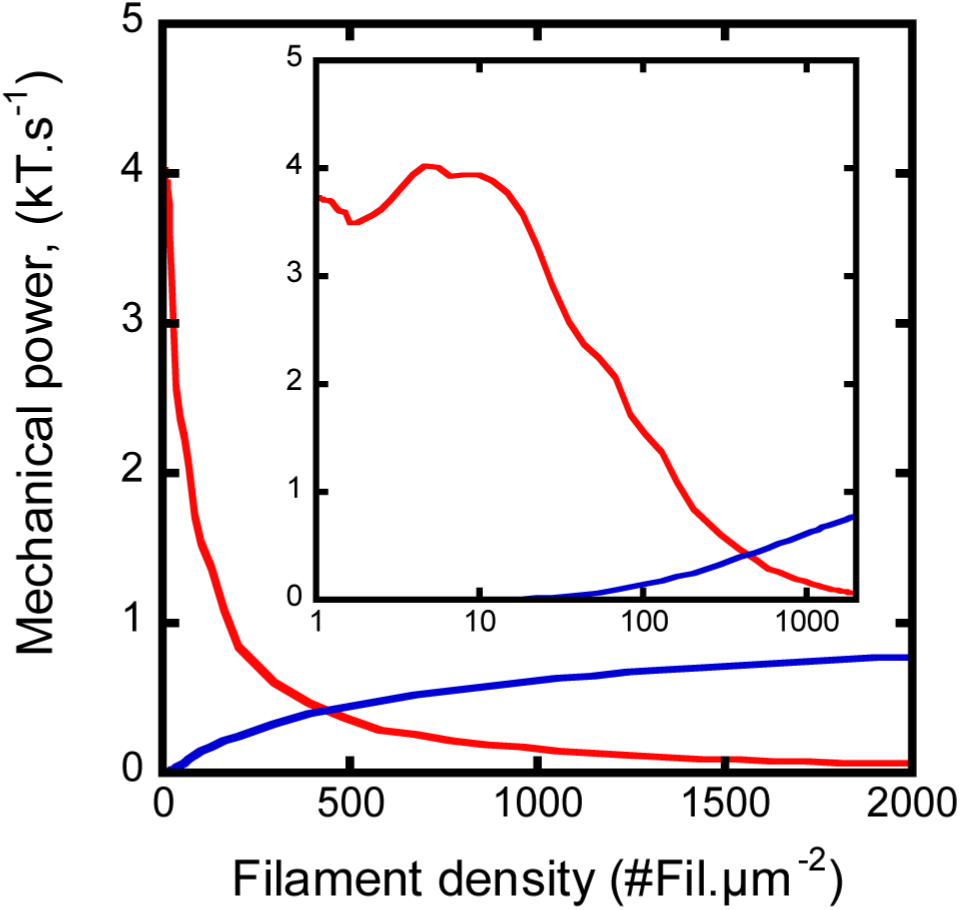
Mechanical power overlap for tethered and non-tethered filaments. Direct comparison of 0° tilt, tethered filaments (red curve) vs. 35° tilt, non tethered filaments (blue curve) displays the complementarity of the two filament architectures.

